# A *Trypanosoma brucei gambiense* gene variant confers human serum resistance

**DOI:** 10.64898/2026.06.16.732631

**Authors:** Nicola Minshall, Sourav Banerjee, Olivia Macleod, Helena Webb, Alex Cook, Matthew Higgins, Mark Carrington

**Affiliations:** Department of Biochemistry, University of Cambridge, Tennis Court Road, Cambridge CB2 1QW, UK; Department of Biochemistry, University of Oxford, South Parks Road, Oxford OX1 3QU, UK; Kavli Institute for Nanoscience Discovery, Dorothy Crowfoot Hodgkin Building, University of Oxford, South Parks Rd, Oxford, OX1 3QU, UK

## Abstract

Most species of African trypanosomes cannot infect humans due to the presence in our blood of trypanolytic factors (TLFs). These lipoprotein particles contain the apolipoprotein L1 (ApoLI) toxin which, when internalised by trypanosomes, forms pores and causes cell death. However, two subspecies of *Trypanosoma brucei* have evolved resistance to TLFs and cause Human African Trypanosomiasis (HAT). The mechanism of resistance of *T. b. rhodesiense* requires a single additional molecule, the serum resistance associated protein SRA. However, the mechanism of resistance of *T. b. gambiense*, which causes HAT in West Africa, has not been fully understood. Here we identify a single polymorphic variant of a PLAC8-domain containing protein which is required for human serum resistance and call this *T. b. gambiense*-specific resistance variant, TgsRV. African trypanosomes are coated with a dense layer of many copies of one member of the variant surface glycoprotein protein family (VSGs). For cells expressing some VSGs, TgsRV is sufficient for human serum resistance, while cells which express other VSGs require a second protein, TgsGP in addition to TgsRV. We therefore complete the identification of the molecular players required for human serum resistance by the African trypanosomes.

## Introduction

Human African Trypanosomiasis results from systemic infection by one of two subspecies of *Trypanosoma brucei.* The trypanosomes initially infect blood and tissue spaces, at a later stage entering the central nervous system where they cause a range of neurological symptoms including the characteristic disruption of the diurnal rhythm [1]. Despite advances in controlling the spread of the disease [2], a poorly characterised human carrier population and a livestock reservoir complicate elimination [3]. Humans and some other primates have evolved specialised innate immunity to most *T. b. brucei* isolates [4–6]. However, human-infective subspecies have evolved countermeasures [5, 6] and here we identify the genes necessary and sufficient for human infectivity in *T. brucei gambiense*, the major causal agent of the disease.

Human innate immunity to *T. b. brucei* is dependent on circulating Trypanolytic Factors (TLF1 and TLF2), which are subsets of high-density lipoprotein (HDL) particles [7–9]. In addition to the HDL core of apolipoprotein A1 and lipid, TLFs contain two primate specific proteins, apolipoprotein L1 (ApoL1) and haptoglobin related protein (Hpr) [10, 11]. The entry of TLF1 into trypanosomes is facilitated by binding of Hpr-haemoglobin complex to the trypanosome haptoglobin-haemoglobin receptor (HpHbR), although absence of the receptor delays, but does not prevent cell death [12]. TLF2 differs from TLF1 as it contains a single additional germline IgM [10] that is proposed to facilitate uptake by binding the variant surface glycoproteins (VSG) which coat trypanosome cell surfaces [13]. After endocytosis, APOL1 forms pH-gated pores in trypanosome membranes resulting in lysosomal swelling [11], loss of integrity of the plasma membrane [14] and cell death.

*T. brucei* has evolved resistance to TLFs at least twice, resulting in human infective trypanosomes. *T. b. rhodesiense* has acquired a toxin-antitoxin system, gaining a single *T. b. rhodesiense* specific protein, serum resistance associated protein (SRA) which binds APOL1 and prevents pore formation [11]. SRA is both necessary and sufficient for resistance to TLFs as loss of SRA expression results in susceptibility and transgenic *T. b. brucei* cell lines which express exogenous SRA can grow in human serum. In *T. b. gambiense*, the mechanism of resistance to TLF has not been fully understood. The *T. gambiense* specific glycoprotein (TgsGP) has been reported as necessary but not sufficient for serum resistance [15, 16]. In addition, *T. b. gambiense* has a point mutation in HpHbR which reduces TLF1 binding affinity and uptake [17, 18], as well as changes in cysteine protease activity [16, 19]. However, these changes are not sufficient to confer the ability on *T. b. brucei* to grow in human serum, highlighting that an essential part of the mechanism of serum resistance remains unknown. Here, we have identified the missing component present in *T. b. gambiense* which, together with TgsGP, is sufficient to confer human serum resistance to *T. b. brucei*.

## Results

### TgsGP is not necessary for human serum resistance by T. b. gambiense

To determine the factors required for human serum resistance of *T. b. gambiense*, we started by deleting the TgsGP gene in the *T. b. gambiense* Eliane cell line by replacing it with a G418 resistance cassette [15] (Figure 1A). Transfected trypanosomes were maintained in media containing G418 and lacking human serum. Individual clones were screened for loss of TgsGP expression by western blotting, identifying nine G418-resistant clones which lack TgsGP (N1-9) and six parental clones expressing TgsGP (P1-6) (Figure 1B). RNA sequencing analysis confirmed the lack of TgsGP in the G418-resistant clones (Supplementary Table 1). These clones expressed a range of different VSGs as seen from variation in mobility of this most abundant protein in cell lysate (Figure 1B).

**Figure 1:**
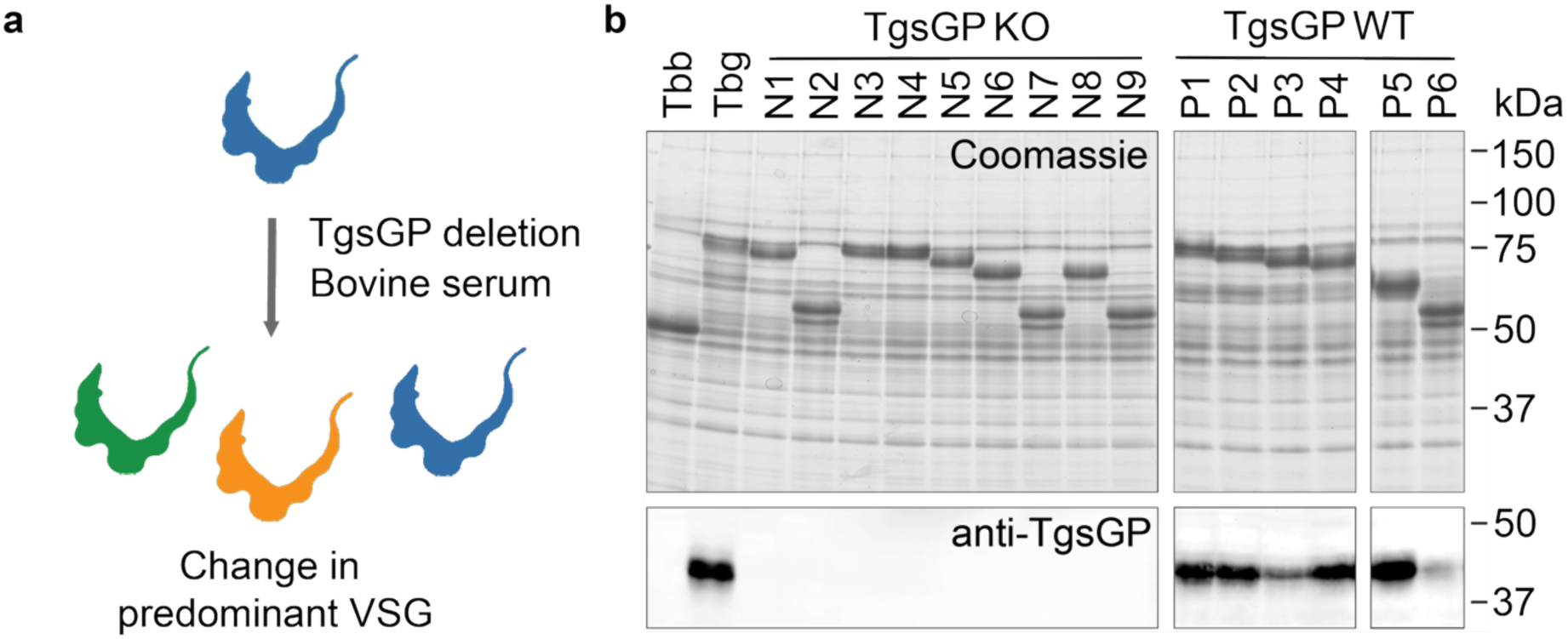
Development of T. b. gambiense isolates lacking TgsGP which express different VSGs. **a.** TgsGP was deleted from a *T. b. gambiense* Eliane cell line and clones were isolated after growth in media containing bovine serum. **b.** Assessment of clones 1-9 (N1-N9) from a *T. b. gambiense* line lacking TgsGP (TgsGP KO) and clones (P1-P6) from a wild-type line (TgsGP WT) expressing TgsGP. The lower panel shows a Western blot probed using an antibody binding to TgsGP. The upper panel shows a Coomassie blue stained gel on which VSG is visible as the most abundant protein in the stained gel, with difference in mobility indicating a difference in VSG identity.

Previous studies reported that TgsGP is necessary for *T. b. gambiense* growth in human serum [15, 16]. To test this, we transferred five TgsGP positive and five TgsGP negative cell lines into media containing 10% human serum and cumulative cell density was determined over 5 days (Figure 2A and B). The human serum used was isolated from two different individual donors and was neither pooled nor heat inactivated. The TgsGP positive clones proliferated in media containing either human or bovine serum as expected, with a 6- to 10-fold increase in cell number over 24 hours (Figure 2B). The TgsGP negative clones proliferated similarly in media containing bovine serum. However, unexpectedly, some of the TgsGP negative clones also proliferated in human serum, with strikingly different growth profiles depending on the identity of the serum donor (Figure 2B). TgsGP negative clone N7 continued to proliferate at a normal rate immediately after transfer to media containing human serum from donor 1. In contrast, the other four TgsGP negative clones (N1, N3, N5, and N6) first decreased in cell density to <1 x 10^4^ cells/ml within a few hours of entering human serum, before starting to proliferate with growth rates which matched those of the TgsGP positive cells. The opposite pattern was observed in human serum from donor 2, with N1, N 3, N5 and N6 continuing to proliferate in human serum, albeit at a reduced rate, while N7 decreased to <1 x 10^4^ cells/ml within a few hours before proliferating cells emerged after 3 days. The proliferation of all TgsGP negative populations emerging after selection in human serum was stable over multiple passages in medium containing the same donor human serum as that used in the original selection, showing that *T. b. gambiense* lines can grow in the presence of human serum despite the absence of TgsGP.

**Figure 2:**
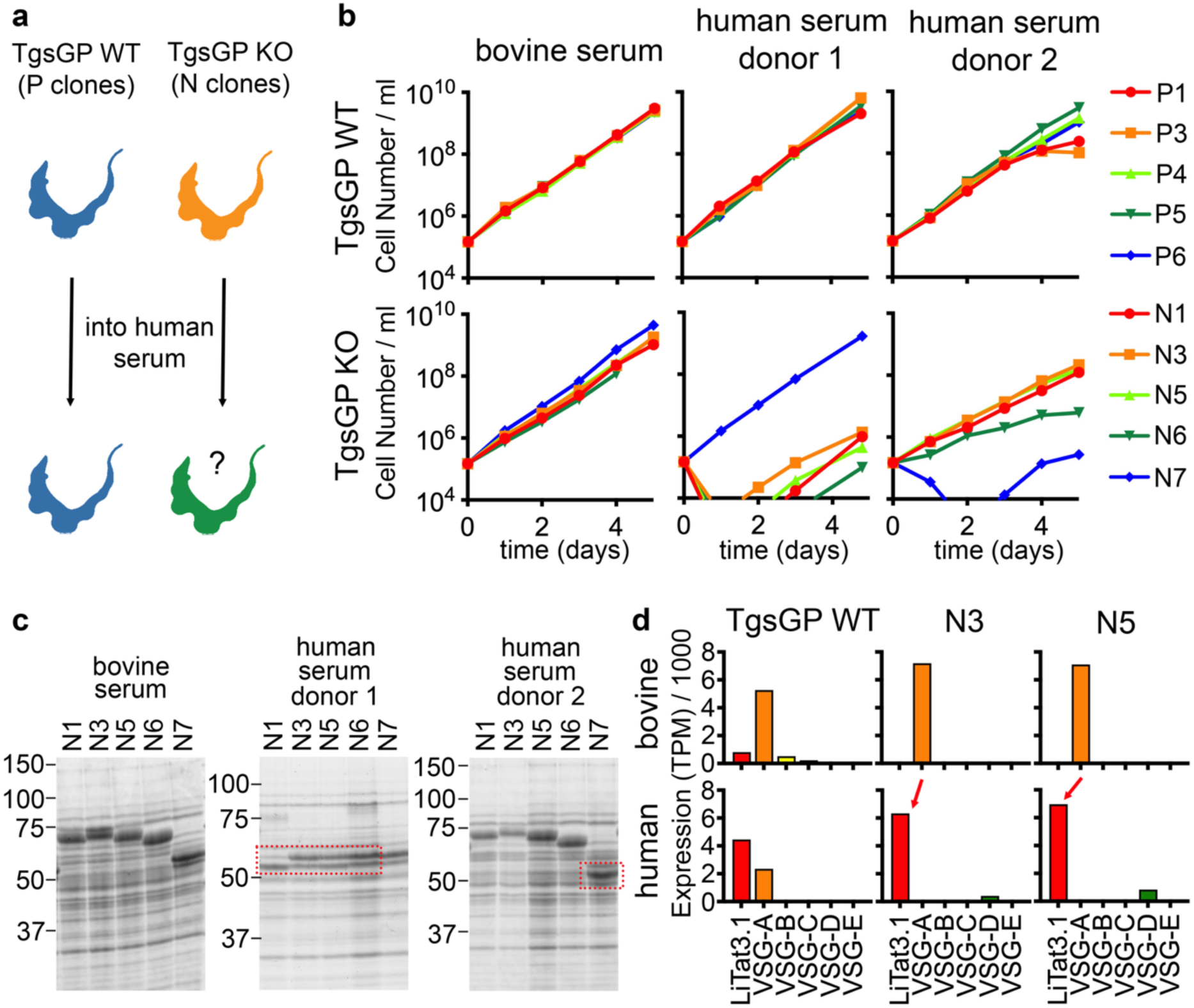
Growth in human serum of T. b. gambiense lacking TgsGP is associated with specific VSGs. **a.** TgsGP was deleted from *T. b. gambiense* and cells were cultured in human serum. **b.** Growth of *T. b. gambiense* clones with (P) and without (N) TgsGP in bovine serum or human serum from two different donors. Cumulative growth curves are shown over 5 days for 5 independent clones. **c.** Changes in VSG expression in cells from (b) analysed after 5 to 10 days. Red dashed boxes show where VSG expression has changed on growth in human serum. **d.** VSG transcript levels determined by RNA sequencing from *T. b. gambiense* or from clones N3 or N5, after growth in bovine or human serum, showing a switch in the predominant VSG in lines N3 and N5 on transfer to human serum.

### Human serum resistance in absence of TgsGP is dependent upon the expressed VSG

To understand the changes which allowed growth in human serum, we first analysed whole cell lysates by gel electrophoresis (Figure 2C). This showed that growth of the five TgsGP negative clones in bovine serum resulted in no change in gel mobility of the expressed VSG. Similarly, clones whose growth was unaffected by the addition of human serum continued to express the same VSG. In contrast, clones which first decreased in cell density in the presence of human serum, before starting to growth after 2-3 days, showed expression of a VSG with a different gel mobility from that in the starting clone. (Figure 2C).

We next analysed the same clones using RNA sequencing to determine whether changes other than the expressed VSG had occurred (Figure 2D, Supplementary Table 1). RNAseq data were used to assemble sequences of expressed VSG cDNAs. Of the six VSGs identified, one was LiTat3.1, previously identified as being favoured during growth in media containing human serum [20], while the others were novel, VSG-A to VSG-E (Figure 2D). The initial clones, grown in bovine serum, contained cells expressing various VSGs. Notably, TgsGP-negative clones 3 and 5, which had adapted to growth in human serum, both changed from predominantly expressing VSG-A to predominantly expressing LiTat3.1, almost certainly due to outgrowth of a small subset in the original population (Figure 2D). As the experiments had been conducted in 10% human serum, we next increased the selection pressure by growing the lines in media containing 50% individual donor human serum (Figure 3). Guided by RNAseq analysis, we selected TgsGP positive (P6) and negative (N7) lines that predominantly expressed LiTat3.1. In serum from donor 1, both clones proliferated at the normal rate (Figure 3A) and RNAseq analysis showed that they continued to express LiTat3.1 (Figure 3B, Supplementary Table 1). In serum from donor 3, TgsGP positive cells proliferated normally whereas the TgsGP negative cells decreased in density to <1×10^4^ cells/ml before a normally growing population emerged 4 days later (Figure 3A) which predominantly expressed VSG-E (Figure 3B, Supplementary Table 1).

**Figure 3:**
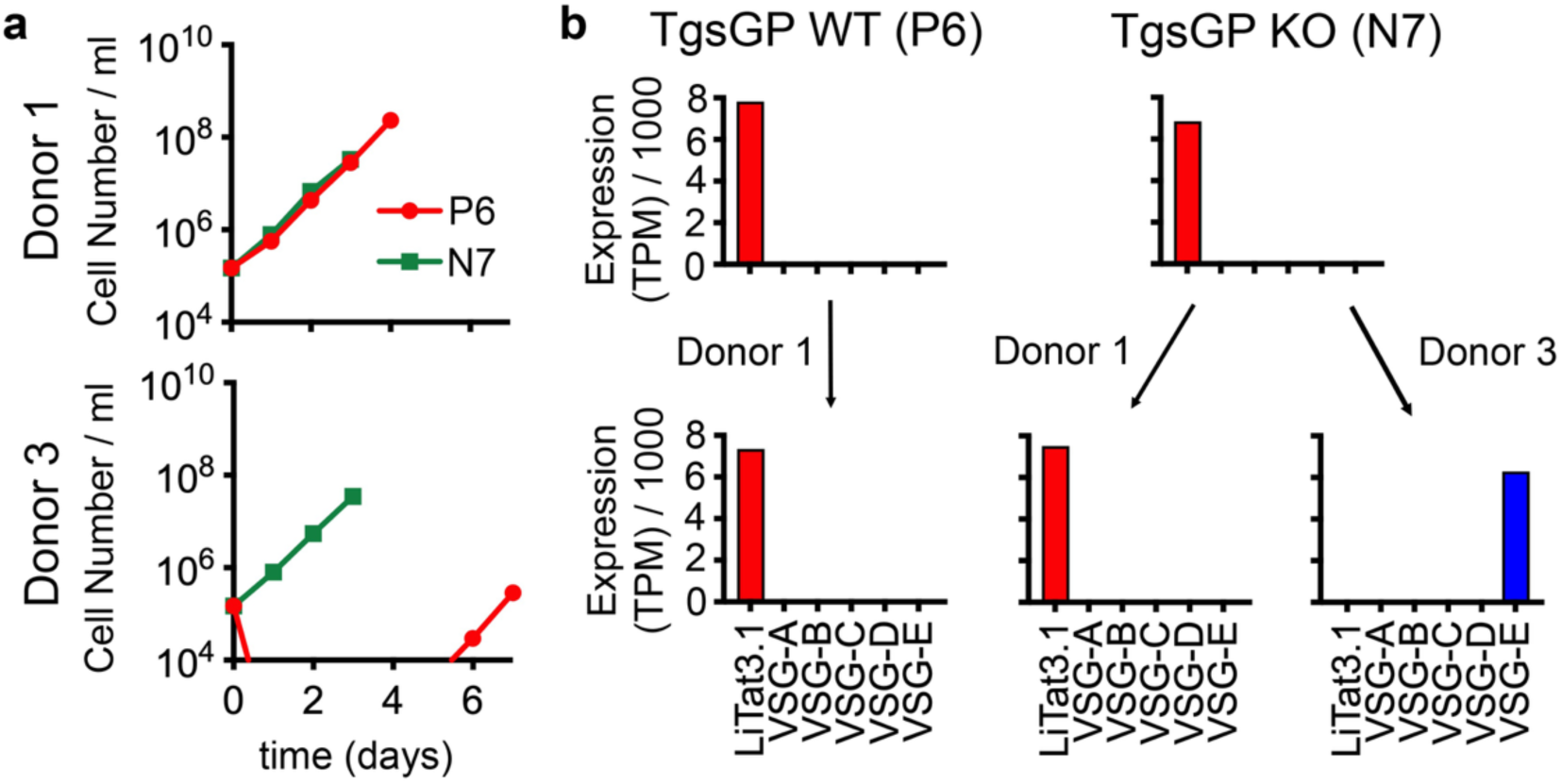
Growth of T. b. gambiense lacking TgsGP depends on the nature of the human serum. **a.** Cumulative growth curves of TgsGP positive (P6) negative (N7) cells expressing VSG LiTat3.1 in 50% human serum from two donors, showing differential growth. **b.** Analysis of VSG transcripts levels by RNA sequencing for cells from a., showing a switch in VSG expression for N7 in serum from donor 3.

These experiments show that TgsGP is not necessary for growth in high concentrations of human serum but that the requirement for TgsGP depends on the nature of the expressed VSG. In the absence of TgsGP, the ability to proliferate is associated with the identity of the expressed VSG, with a gain of growth in human serum associated with a change in the predominant VSG expressed by the population. This is consistent with a model in which interactions between a variable component in the serum and VSGs determines killing or survival. When the trypanosomes express a susceptible VSG, TgsGP is then required for growth.

### Identification of T. b. gambiense specific polymorphisms required for human serum resistance

While previous studies have reported factors which are required for *T. b. gambiense* to retain resistance to human serum [15, 16], no study has so far identified a set of *T. b. gambiense* factors which can confer human serum resistance when introduced into *T. b. brucei*. To identify such factors, we used a gain of function screen in which fragments of *T. b. gambiense* genomic DNA were introduced into a human serum sensitive *T. b. brucei* line (Figure 4). To generate a recipient cell line, we started with *T. b. brucei* Lister 427 and introduced three changes previously associated with human serum resistance in *T. b. gambiense* (Figure 4A). First, both alleles of the HpHbR gene were replaced with puromycin and hygromycin resistance cassettes (Supplementary Figure 1). Second, a TgsGP transgene was inserted in the tubulin locus along with a G418 resistance cassette and we selected a clone with TgsGP expression similar to that of *T. b. gambiense* (Supplementary Figure 1). Third, the expressed VSG2 was replaced by LiTat3.1 which was favoured in human serum resistant *T. b. gambiense* grown in culture (Supplementary Figure 1) [20]. When cultured with 5% pooled human serum, the cells were killed with no resistant population emerging, confirming that these changes are not sufficient for human serum resistance of *T. b. gambiense*.

**Figure 4:**
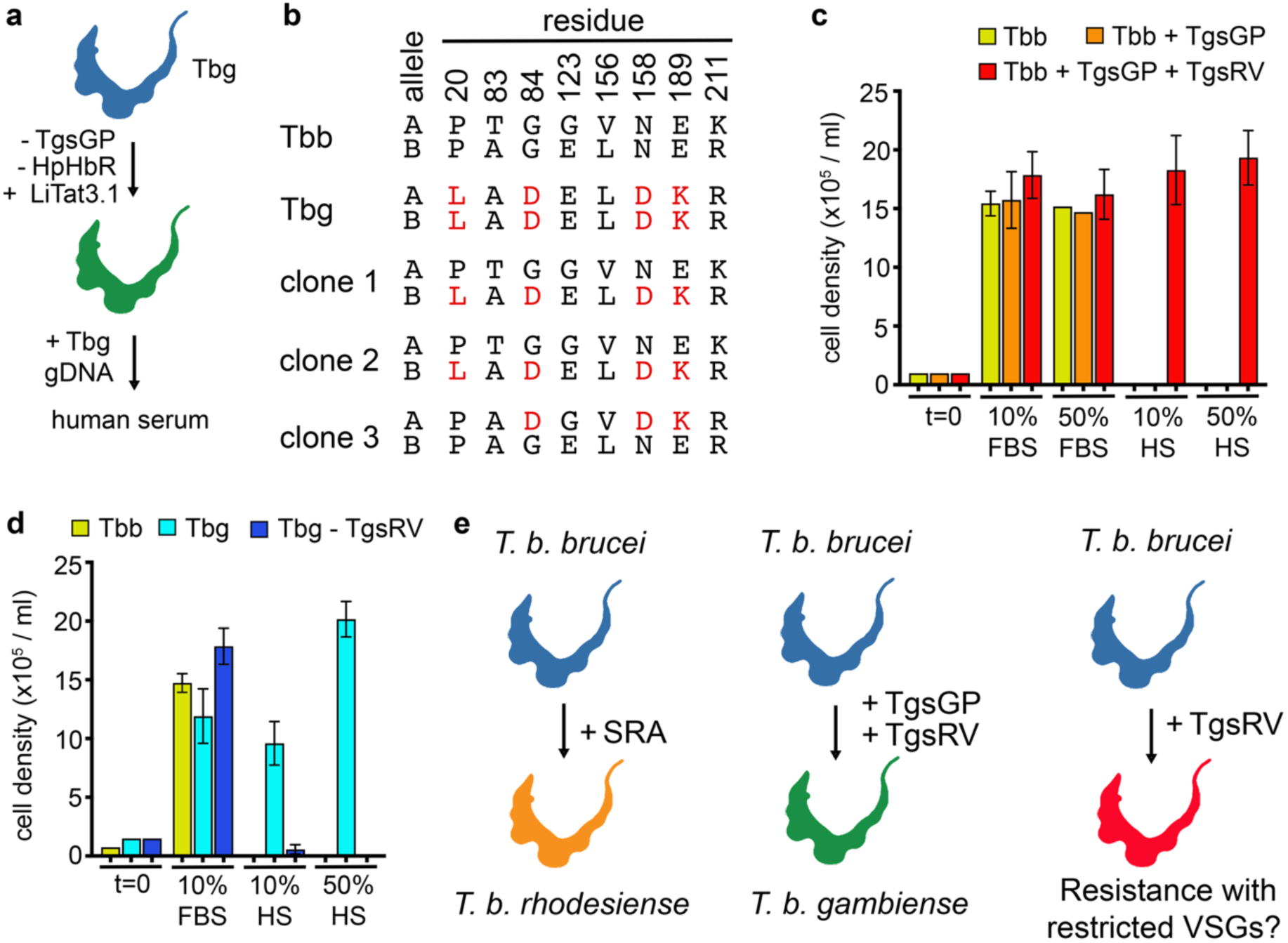
Identification of TgsRV as necessary for growth in T. b. gambiense in human serum. **a.** *T. b. brucei* lacking HpHbR and expressing TgsGP and VSG LiTat3.1 was transfected with fragmented *T. b. gambiense* genomic DNA and selected for growth in human serum. **b.** The identity of specific polymorphic residues found in the two alleles (A and B) of the equivalent genes, Tb427_020016000 (Tbb) and Tbg.972.2.1820 (Tbg) and in the three cell lines selected in human serum (clones 1 to 3). **c.** Cell density after growth for 24 hours of *T. b. brucei*, or *T. b. brucei* lines expressing just TgsGP or both TgsGP and TgsRV in 10% and 50% fetal bovine serum (FBS) or pooled human serum (HS). Averages are plotted and standard deviations shown are from at least 5 technical replicates of Tbb and Tbb+TgsGP cell lines along with 7 biological replicates ofTbb+TgsGP+TgsRV. **d.** Cell density after growth for 24 hours of *T. b. brucei* (Tbb)*, T. b. gambiense* (Tbg) or a *T. b. gambiense* line lacking TgsRV (Tbg – TgsRV) in 10% and 50% fetal bovine serum (FBS) or pooled human serum (HS), two distinct pools of human serum were used. Averages and standard deviations are shown, for two Tbg clones, three Tbg-TgsRV clones or Tbb clone replicates. Starting cell densities were 7.5 x 10^4^ cells/ml for *T. b. brucei* cells and 1.5 x 10^5^ cells/ml for *T. b. gambiense* cell lines. **e.** The essential determinants of human serum resistance in *T. b. rhodesiense* (SRA) and *T. b. gambiense* (TgsGP and TgsRV) with TgsRV alone providing human serum resistance in a VSG restricted manner.

We next introduced randomly sheared genomic DNA from *T. b. gambiense* Eliane into this reporter line and initiated selection after 6 h by addition of 5% pooled human serum and splitting transfectants into 298 flasks (Figure 4A). After 5 days, three flasks contained rapidly proliferating cells. The reporter line and resistant clones were analysed by RNAseq with sequence data mapped for differential expression against *T. b. brucei* Lister427 and *T. b. gambiense* DAL972 genomes. TgsGP and VSG expression were assessed using RNAseq analysis (Supplementary Table 2) and Western blotting (Supplementary Figure 2A) and no changes were detected. Instead, differential expression analysis revealed all three human serum resistant clones to contain genetic changes in a single gene labelled Tbg.972.2.1820 in *T. b. gambiense* DAL972 and Tb427_020016000 in *T. b. brucei* Lister427 (Supplementary Table 2). A comparison of the sequences of Tbg.972.2.1820 and Tb427_020016000 in the reference genomes showed 8 polymorphic residues (Supplementary Figure 2B). The *T. b. brucei* reporter line contained two polymorphic alleles, whereas the *T. b. gambiense* Eliane alleles were homozygous (Figure 4B). Four polymorphisms present in Tbg.972.2.1820 (L20, D86, D158 and K189) are absent in Tb427_020016000 (Figure 4B). In resistant clones 1 and 2, the B allele included all four *T. b. gambiense* specific residues, whereas in resistant clone 3, the A allele includes three, D86, D158 and K189, demonstrating a dominant effect (Figure 4B). We named Tbg.972.2.1820 the *T. b. gambiense* specific resistance variant, TgsRV.

To validate these finding, we started with the standard laboratory cell line *T. b. brucei* L427 cell lines expressing VSG2 which is readily killed by human serum. These were transfected with either TgsRV alone or both TgsGP and TgsRV and were grown in pooled human serum (Figure 4C). While the starting cell line and cells expressing only TgsGP did not proliferate, the line expressing both TgsGP and TgsRV did proliferate. Seven clones were selected for analysis and the sequences of both alleles of TgsRV were determined after further growth in human serum. All had incorporated at three of the polymorphic residues from TgsRV (D86, D158 and K189) into one of the two alleles (Supplementary Figure 3). The proliferation of these cell lines in culture was measured over 24 h (Figure 4C) and the ∼16-fold increase in cell number over 24 h in both 10% and 50% human serum is similar to its growth in the absence of human serum and exceeds *T. b. gambiense* growth in the same medium (Figure 2B). This observation indicated that expression of both TgsGP and TgsRV is sufficient to confer robust resistance to human serum in the VSG2 background.

We next tested whether TgsRV is necessary for proliferation of *T. b. gambiense* in pooled human serum using targeted gene deletion. The two alleles of TgsRV were replaced with blasticidin and G418 resistance cassettes (Supplementary Figure 4). Three clones were selected by resistance to both antibiotics in absence of human serum and deletion was confirmed first by PCR (Supplementary Figure 4) and short read genome sequencing (Supplementary Table 3). Proliferation of these three clones over 24 h was compared with that of two clones of *T. b. gambiense* Eliane and *T. b. brucei* L427 (Figure 4D). While all cell lines proliferated in the absence of human serum, *T. b. brucei* L427 was killed in media containing human serum whereas *T. b. gambiense* proliferated. The three TgsRV deletion clones all failed to proliferate in 10% human serum with a reduced number of cells visible after 24 h. However, all three deletion clones were killed by 50% human serum and no resistant cells emerged over 7 days (Figure 4D).

We next checked whether the polymorphisms found in TgsRV were common across all *T. b. gambiense* isolates. The sequences of TgsRV were extracted from the assembled reference genome (Daloa 972) and 11 other isolates from short read archives (Supplementary Table 4). All isolates were homozygous for the three polymorphisms, D86, D158 and K189, consistent with a central role in TLF resistance in *T. b. gambiense*. In contrast, the polymorphisms were not present in the genomes of 7 isolates of *T. b. rhodesiense*, the human infective form present in East Africa (Extended Data 1).

## Discussion

These findings answer the long-standing question about what makes *T. b. gambiense*, the main causal agent of Human African Trypanosomiasis, human serum resistant. Our main finding is to identify TgsRV as the one essential determinant of human serum resistance, as deletion of TgsRV renders *T. b. gambiense* sensitive to human serum and the introduction of TgsRV into *T. b. brucei* enables it to proliferate in human serum in the presence of TgsGP.

TgsRV encodes a 215 residue protein of unknown function with a PLAC8 domain between residues 59 to 206, with the remainder predicted to be disordered. PLAC8 domain-containing proteins are found in a wide range of eukaryotes, often associated with membranes, but with no obvious common function [21]. The *T. b. brucei* homologue of TgsRV is localised to the endosomal compartment of procyclic trypanosomes [22] and we confirmed this in cultured bloodstream form *T. b. brucei* and *T. b. gambiense* (Supplementary Figure 5). This places TgsRV in a location in which it could potentially interact directly with TLF particles and/or ApoLI. Indeed, TgsRV is dominant in generating human serum resistance (Figure 4B), suggesting that the three changes from the *T. b. brucei* homologue might generate a binding site which directly or indirectly prevent the activation or action of the ApoL1 pore preventing trypanosome cell death.

Our experiments also reveal more about the role of TgsGP. TgsRV and TgsGP are together sufficient to provide human serum resistance in all cases tested in this study. However, whether TgsGP is required depends on the nature of the expressed VSG. *T. b. gambiense* clones can proliferate in the absence of TgsGP when they express some VSGs and, when the VSG is not compatible with resistance, a change in VSG is sufficient to allow a gain of growth in human serum in the absence of TgsGP (Figure 2). Also striking is that human serum from different donors is compatible with different VSGs, suggesting that this effect is due to interactions between VSGs and a variant factor presence in human serum. While further studies are required to establish the nature of this variant factor, one possibility is that it is the IgM core of the TLF2 particles, perhaps with differential interactions between VSGs and IgM affecting TLF2 uptake and therefore the requirement for TgsGP to inactivate TLFs or ApoLI. It is likely that TgsGP is retained in *T. b. gambiense* to allow it to retain human serum resistance while maintaining the ability to use the full repertoire of VSGs.

This discovery that two factors, TgsGP and TgsRV are sufficient to confer human serum resistance to *T. b. brucei*, completes the discovery of the factors required for *T. b. gambiense* to survive in human serum and open the way to discover their mechanism of action.

## Methods

### Cell lines and culture conditions

*T. b gambiense* Eliane was obtained from Annette MacLeod (University of Glasgow). *T. b. brucei* Lister 427 expressing VSG2 was obtained from Mike Ferguson (University of Dundee) and adapted to culture. All cell lines were maintained in HMI11 medium supplemented with heat inactivated foetal bovine serum and/or human serum [23] and were maintained in log phase between 1×10^5^ and 2×10^6^ /ml. Human serum was obtained from individual donors; pooled human serum for the large scale screen was a 1:1 mixture of commercial (Appleton Woods CSR130) and pooled serum from donors. Pooled serum for growth experiments were mixtures from individual donors. Cell density was determined using a haemocytometer.

### Protein gels and western blotting

Standard methods were used for SDS-polyacrylamide gels and western blotting. Rabbit anti-TgsGP was raised against recombinant protein corresponding to residues 21 to 389 with a C-terminal His tag. The protein was expressed in CHO cells and purified by affinity chromatography and size exclusion chromatography and the antiserum produced by Covalab. Anti-TgsGP was used at 1:5000. Anti-221, 1:5000 [24] and anti-SCD6, 1:10000 [25].

### RNA and DNA preps

RNA was prepared using Qiagen RNAeasy kits. Genomic DNA for sequencing or for use as PCR templates was prepared using Qiagen Blood and Tissue DNA kits. *T. b. gambiense* DNA for electroporation was prepared using a modified protocol from Wendy Gibson (University of Bristol). Cells were resuspended in 10mM Tris-HCl, 100mM NaCl, 100mM EDTA, pH 8 at 1×10^9^ /ml and 1/9 volume of 10% sodium lauroyl sarcosinate was added to lyse the cells. Proteinase K was added to 100 µg/ml and incubated at 50°C to 4 h. The mixture was phenol/chloroform/isoamyl alcohol extracted and DNA precipitated by adding sodium acetate to 300 mM and 2.5 volumes of ethanol. The genomic DNA was recovered by spooling and dissolved in 10mM Tris-HCl, 0.1mM EDTA, pH 8. DNAse-free RNAse was added to 20 µg/ml and incubated at 37°C for 30 mins. Sodium dodecyl sulphate was added to 0.1% and the proteinase K digestion and phenol/chloroform/isoamyl alcohol extractions were repeated. The genomic DNA was recovered by isopropanol precipitation and spooling and dissolved in 10 mM Tris-HCl, 0.1 mM EDTA pH 8.0

### Transfection and selection

Transgenic trypanosomes were produced using standard procedures with an Amaxa Nucleofector using program X-001. Selection was added 6 h after electroporation. For the gain of function screen, purified *T. b. gambiense* genomic DNA was sheared to ∼1.5 kbp and was introduced into the reporter cell line in 21 separate electroporations, each with 3 x 10^7^ cells and 40 μg genomic DNA. After 6 h the cells were diluted to 2 x 10^5^ /ml and pooled human serum added to 5%. This yielded 298 flasks, each containing 10 ml culture.

### Plasmids, PCR and RT-PCR

Plasmid manipulation and PCR reactions used standard procedures. Details and oligonucleotides sequences are in Supplementary Table 5, and plasmid sequences are in Supplementary Figure 9. For RT-PCR, 300 ng total RNA was incubated at 65°C for 10 mins in a volume of 13 µl containing 0.8 µM T15VN and 80 µM N6 primers. The mixture was snap cooled on ice for 1 min and was then used as the substrate for a 20 µl Transcriptor (Roche) reaction containing 20 U Ribolock RNase Inhibitor (ThermoFisher), 1 mM dNTPs and 10 U Transcriptor Reverse Transcriptase. The reaction was incubated at 25°C for 10 min, 55°C for 30 min and 85°C for 5 min. 0.5 µl of this reaction was used as substrate in a standard PCR.

### Sequencing

Sanger sequencing of plasmids was performed in the Sequencing Facility in the Department of Biochemistry, University of Cambridge. RNAseq was performed by the Beijing Genomics Institute using DNBSEQ Eukaryotic Strand-specific Transcriptome PE150 Resequencing. Short read genome sequencing used Project Information DNBSEQ Animal and Plant WGRS PE100.

### RNAseq and DNAseq analysis pipelines

All analysis was performed using the Galaxy server [26]. Sequence data were run through Trimmomatic [27] with the following parameters: LEADING, 10; TRAILING, 10; SLIDING WINDOW, 5, 15; MINLEN, 50. Paired end reads were then formatted with FASTQgroomer [28]. To identify expressed VSGs from RNAseq data, the sequences were mapped to the *T. b. gambiense* DAL972 genome v68 downloaded from TriTrypDB [29] using Bowtie2 [30] with default settings and unaligned reads were written to a file. The unaligned reads were assembled using Trinity [31] and default settings. Open reading frames of >500 nucleotides were identified using GetORF [32] and mapped to the .gff file using Exonerate. Total reads from above were then remapped to the genome and Trinity assembly combined using Bowtie2. TPM were calculated using FeatureCounts [33] and the bam file. All sequences with a TPM value of >2000 were translated and VSGs identified by amino acid sequence. For the differential expression analysis in the gain of function screen, reads were initially mapped using Bowtie 2 to a genome file made by combining: *T. b. gambiense* Dal972 genome v68, *T. b. brucei* Lister 427 2018 genome v68, and sequences encoding VSG LiTat3.1 and TgsGP. FeatureCounts was used to identify differentially expressed genes. Short read genome sequence data was used to confirm deletion of the Tbg.972.2.1820 gene. Reads were mapped to the *T. b. gambiense* Dal972 genome v68 using Bowtie2. Mosdepth [34] was used to calculate the average read depth from the bam file. Read depth for individual genes was calculated using Mosdepth with regions defined by a .bed file generated from the *T. b. gambiense* Dal972 v68 .gff file using GFF-to-BED [26]. Assembly of Tbg.972.2.1820 homologues from published gDNA sequencing files was performed using Shovill [26] on FASTQ groomer files to assemble contigs.

### mNG tagging and microscopy

Cell lines were modified using *in vitro* assembled Cas9 RNPs [35] to target the tag repair to the relevant sequences [36]. Cells were visualised using epifluorescence microscopy immediately after fixation in 3% formaldehyde .

## Data availability

ENA project accession number is PRJEB107749 and the individual accession numbers are in Supplementary Table 6

## Acknowledgements

This work was funded by a Wellcome Investigator award (217138/Z/19/Z) to MC and MKH. We would like to thank: Annette MacLeod for discussions, the *T. b. gambiense* Eliane cell line and the plasmid construct used to delete the TgsGP gene, Dawn Ahi for phlebotomy and the Cambridge Department of Biochemistry DNA Sequencing Facility.

## Supplementary Figures

**Supplementary Figure 1:**
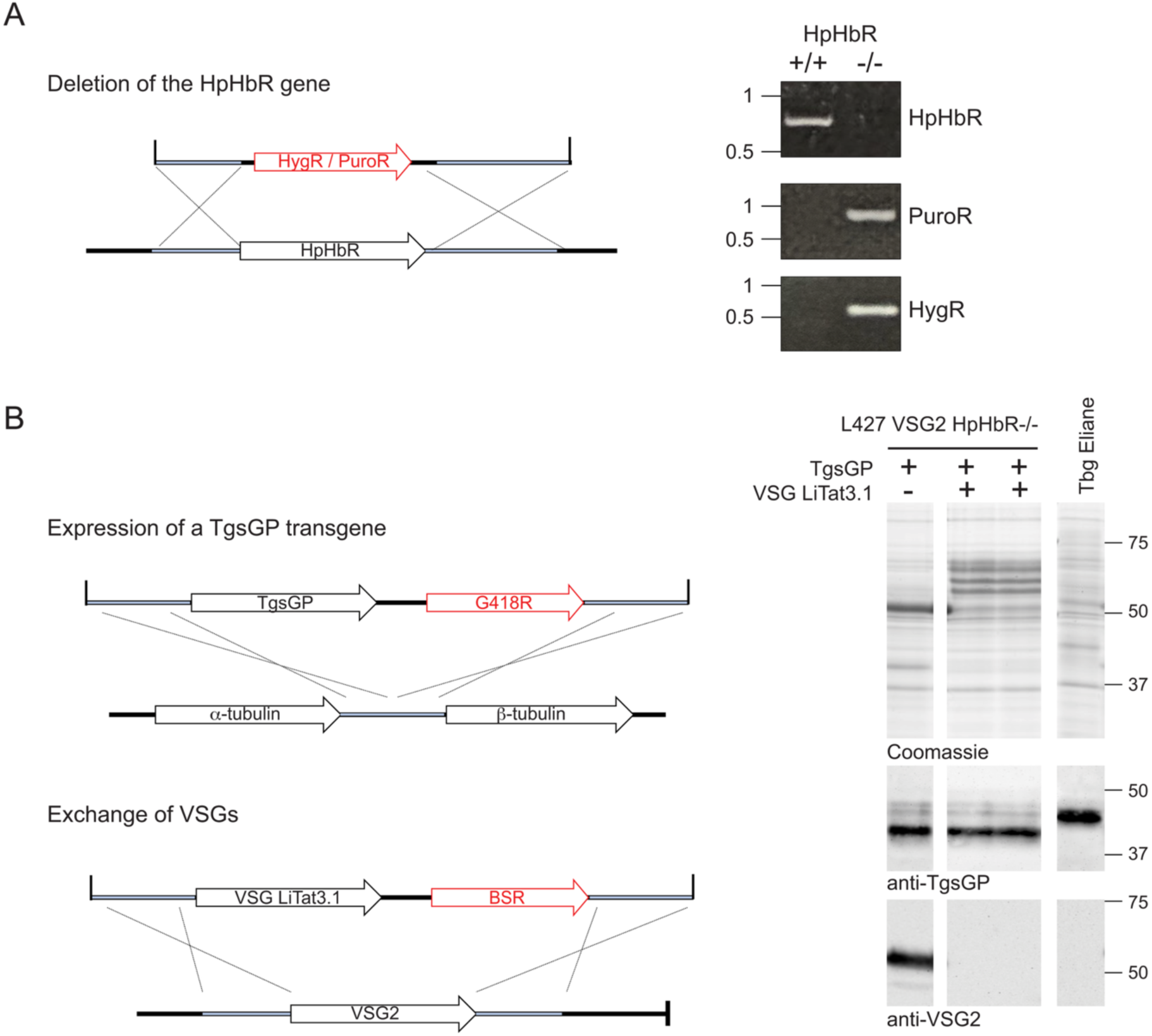
strategy used to generate transgenic T. b. brucei. **a.** Schematics of the strategy used to delete the HpHbR gene and PCR to screen for successful deletion, see Supplementary Table 5 for details of the PCR. **b.** Schematics of the strategies used to insert a TgsGP transgene and replace VSG2 with LiTat3.1. Screening for a successful outcome was by analysis of whole cell lysates and western blotting. Note, processing of N-linked glycans in the Lister 427 parental strain is different to *T. b. gambiense* and results in different mobilities of the same polypeptide.

**Supplementary Figure 2:**
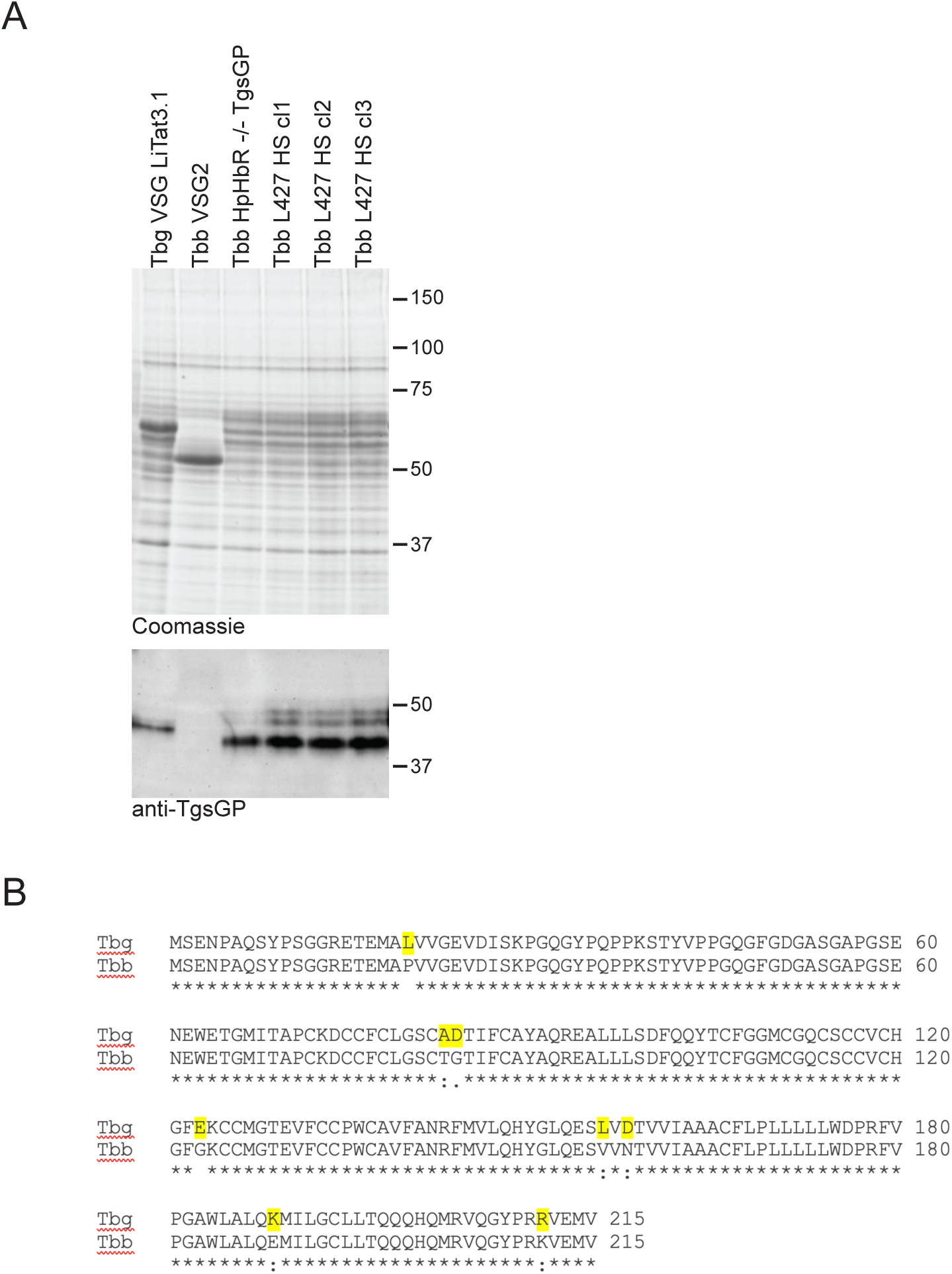
Analysis of cell lines selected in human serum after transfection with T. b. gambiense genomic DNA. A. Analysis of VSG and TgsGP expression in the reporter cell line, Tbb HpHbR-/- TgsGP LiTat3.1, and human serum resistant cell lines Tbb L427 HSR Cl1 to 3, selected in the screen. The VSG is visible in the image of the Coomassie stained gel of whole cell lysate and TgsGP was visualised by western blotting. Note, processing of N-linked glycans in the Lister 427 parental strain is different to *T. b. gambiense* and results in different mobilities of the same polypeptide. B. Comparison of the amino acid sequences of Tbg.972.2.1820 and Tb427_020016000 from the reference genomes. Differences are highlighted in yellow.

**Supplementary Figure 3:**
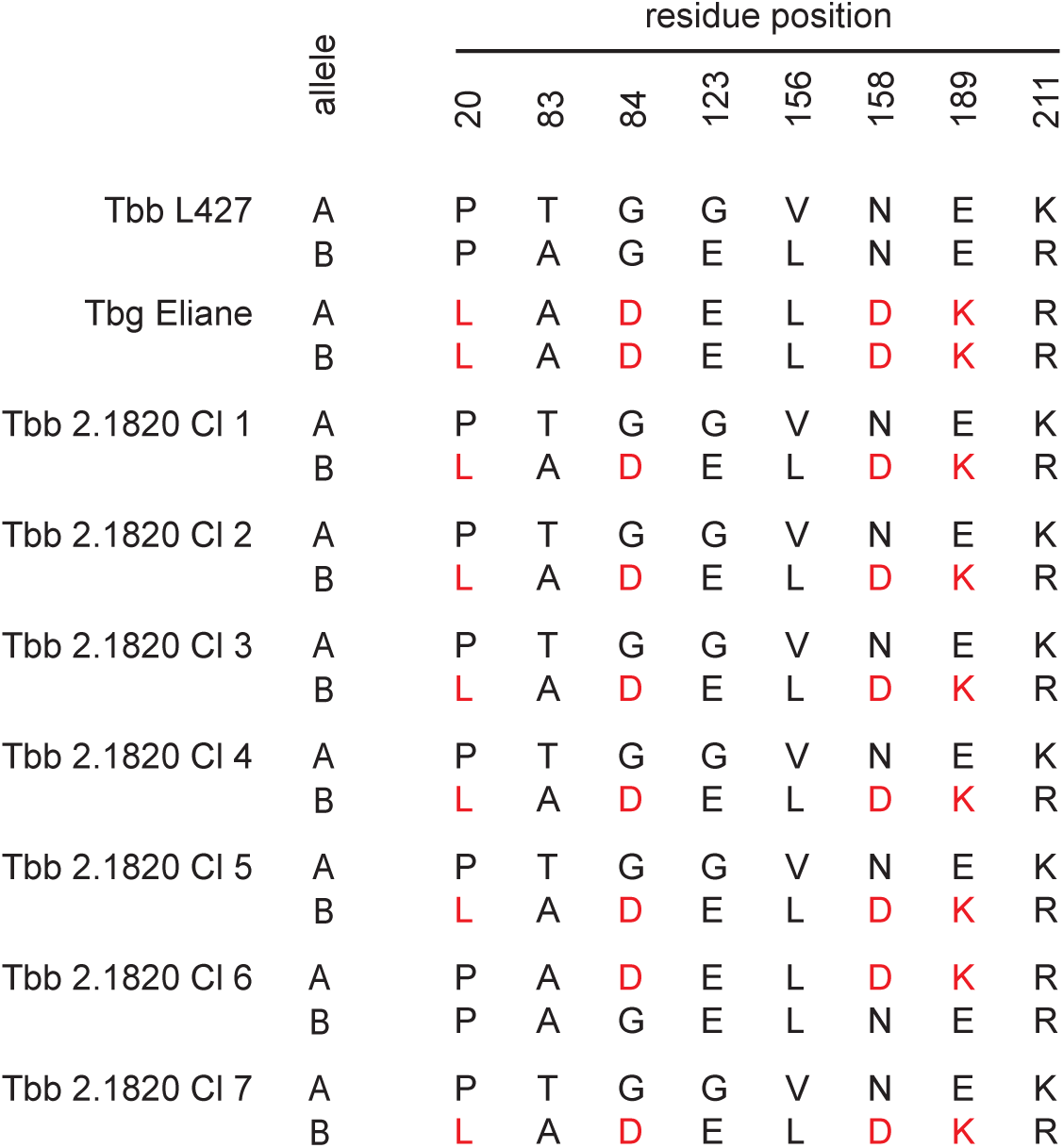
Growth of T. b. brucei expressing TgsGP in pooled human serum is dependent on introduction of TgsRV polymorphisms. Sequence changes in T. *b. brucei* Tb427_020016000 after transfection with TgsRV and selection for growth in human serum

**Supplementary Figure 4:**
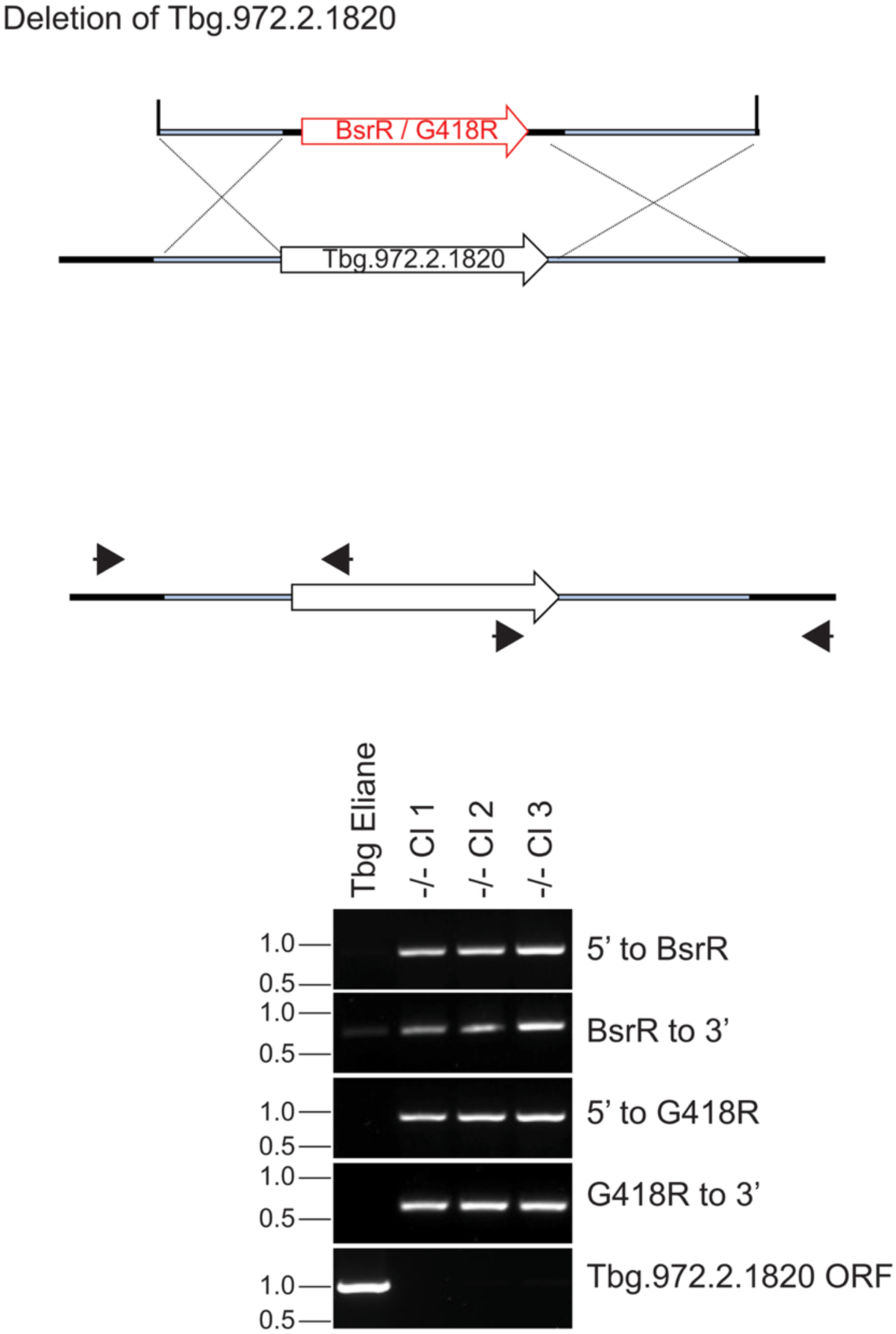
Deletion of the Tbg.972.2.1820 gene. Below the schematic is show output of PCR to screen for successful deletion.

**Supplementary Figure 5:**
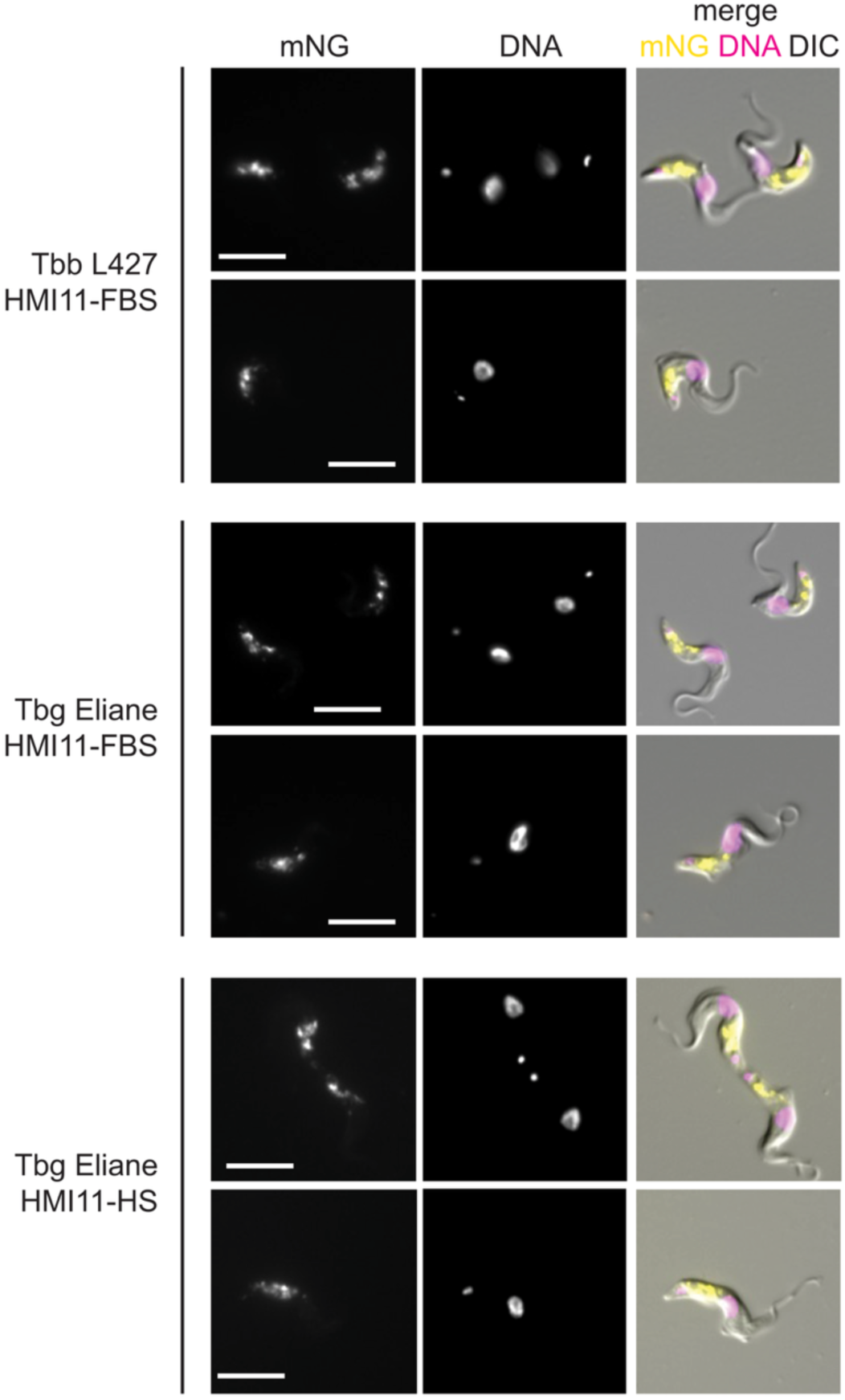
localisation of TgsRV and its T. b. brucei homologue. Subcellular localisation of mNeonGreen tagged Tbg.972.2.1820 and Tb427_020016000 Genes were tagged at the endogenous locus with mNeonGreen (mNG). *T. b. gambiense* clones were observed in both bovine and human serum. The tagged protein localises to endocytic compartments and there are no readily observable differences in either protein quantity or localisation between *T. b. brucei* and *T. b. gambiense* variants or in sera from different organisms.

**Supplementary Table 1.**
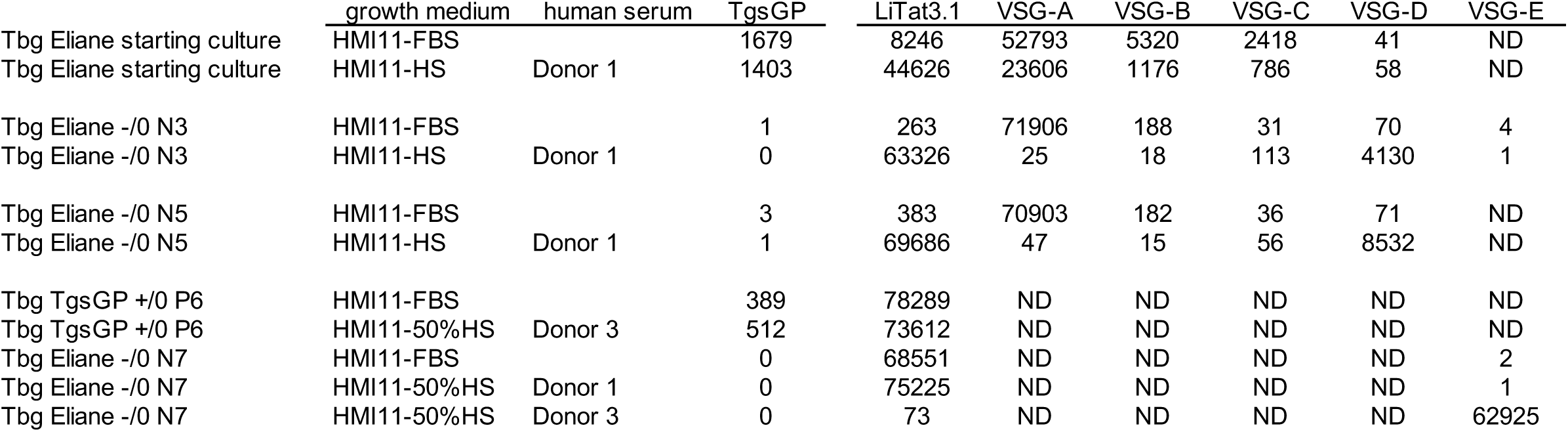
Analysis of RNAseq data to determine TgsGP and VSG mRNA levels in TgsGP WT and null cell lines in FBS and after transfer to HS. Expression level is in TPM. ND: mRNA not detected

**Supplementary Table 2:** Analysis of RNAseq data to determine mRNA levels for TgsGP, VSG, and Tbg.972.2.1820 variants in the gain of function screen.

Expression level is in TPM
|  | Tb427_02<br>0016000<br>allele A | Tb427_02<br>0016000<br>allele B | Tbg.972.<br>2.1820 | TgsGP | VSG<br>LiTat3.1 |
| --- | --- | --- | --- | --- | --- |
| Tbb L427 HpHbR<br>-/- TgsGP LiTat3.1 | 74 | 58 | 4 | 1604 | 50220 |
| Tbb HSR CI1 | 65 | 8 | 55 | 1378 | 55987 |
| Tbb HSR CI2 | 61 | 8 | 50 | 1608 | 56037 |
| Tbb HSR CI3 | 47 | 72 | 64 | 1680 | 56360 |

**Supplementary Table 3:** Analysis of data from genome sequencing before and after Tbg.972.2.1820 deletion.

|  | Read<br>depth |  |
| --- | --- | --- |
|  | Mean | Tbg.972.2.1820 |
| Tbg Eliane | 83.5 | 77.4 |
| Tbg Eliane Tbg.972.2.1820 -/- CI1 | 88.5 | 0.4 |
| Tbg Eliane Tbg.972.2.1820 -/- CI2 | 88.3 | 0.4 |
| Tbg Eliane Tbg.972.2.1820 -/- CI3 | 88.4 | 0.4 |

**Supplementary Table 4.** Accession numbers for Short Read Archive genome sequences.

| Isolate name | Year | Country | Locality | Accession number | Study accession number |
| --- | --- | --- | --- | --- | --- |
| *T. b. gambiense* |  |  |  |  |  |
| Eliane | 1952 | Côte d'Ivoire | Unknown | ERR270826 | PRJEB2486 |
| Daloa972 | 1986 | Côte d'Ivoire | Daloa | FN554965 | PRJEA40697 |
| Abtor | 1983 | Côte d'Ivoire | Vavoua | ERR270831 | PRJEB2486 |
| B4_4163P | 2004 | Côte d'Ivoire | Bonon | ERR270859 | PRJEB2486 |
| B7_2 | 2001 | Côte d'Ivoire | Oumé | ERR270869 | PRJEB2486 |
| Yad | 1998 | Côte d'Ivoire | Sinfra | ERR270892 | PRJEB2486 |
| T21_4A | 1999 | Côte d'Ivoire | Grand-Zathry | ERR270891 | PRJEB2486 |
| Mos | 1974 | Cameroon | Mbam | ERR270829 | PRJEB2486 |
| Kide | 1983 | Côte d'Ivoire | Gagnoa | ERR270836 | PRJEB2486 |
| Dob112_Kivi | 2002 | Guinea | Boffa | ERR270858 | PRJEB2486 |
| Deola | 1984 | Côte d'Ivoire | Daloa | ERR270839 | PRJEB2486 |
| Bim_1 | 1975 | Cameroon | Campo | ERR270838 | PRJEB2486 |
| B13_9 | 1998 | Guinea | Dubreka | ERR270886 | PRJEB2486 |
| *T. b. rhodesiense* |  |  |  |  |  |
| EATRO2340 | 1977 | Kenya | Unknown | ERR019097 | PRJEB2167 |
| EATRO3 | 1960 | Uganda | Unknown | ERR019098 | PRJEB2167 |
| GYBO | 1984 | Tanzania | Tanzania | ERR270817 | PRJEB2486 |
| K975 | 1970 | Kenya | Kiboko | ERR270820 | PRJEB2486 |
| Z210 | 1982 | Zambia | Luangwa | ERR270821 | PRJEB2486 |
| N1052 | 1961 | Kenya | Nyanza | ERR270825 | PRJEB2486 |

**Supplementary Table 6:**
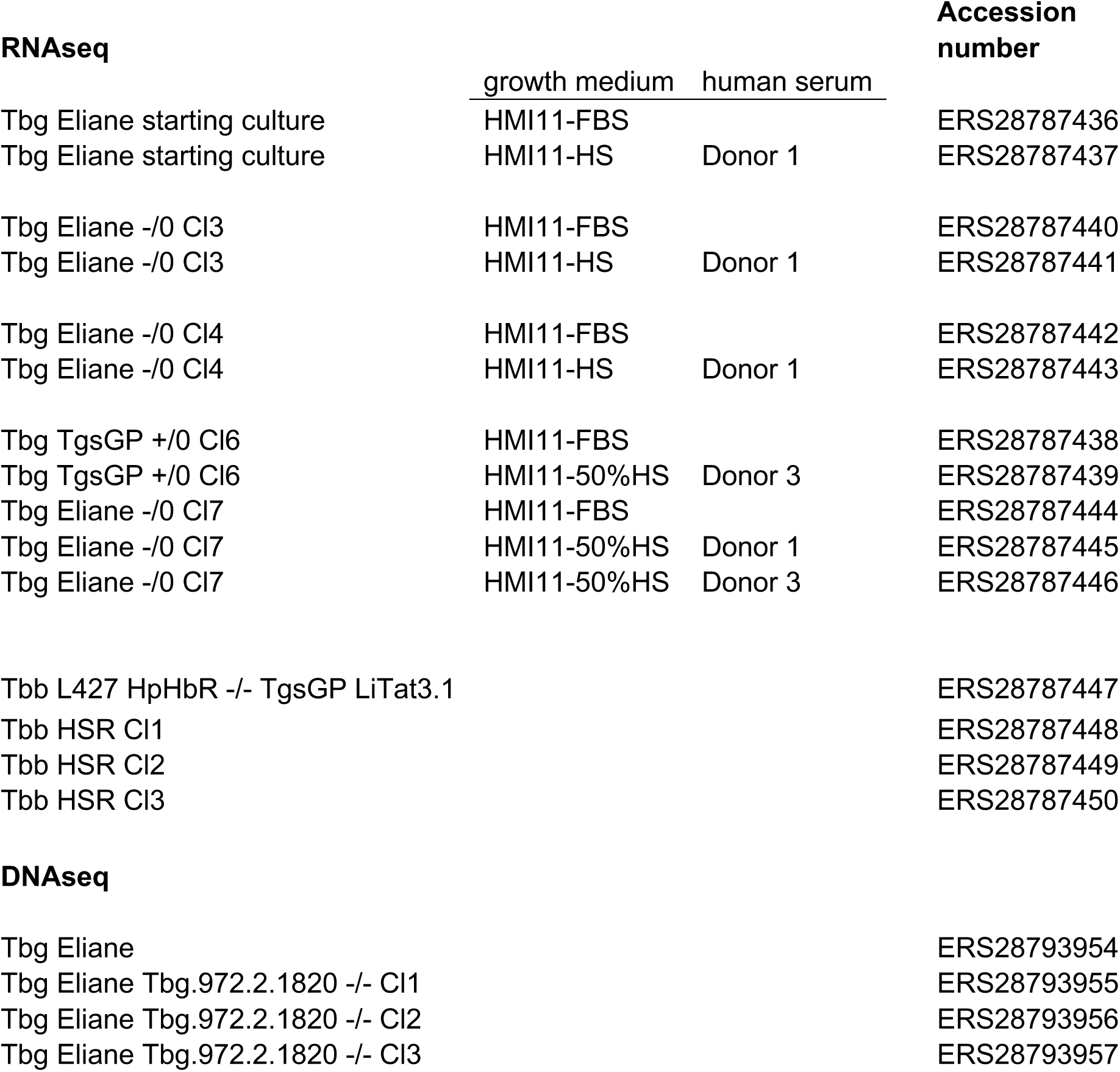
Accession numbers.

**Extended Data 1**

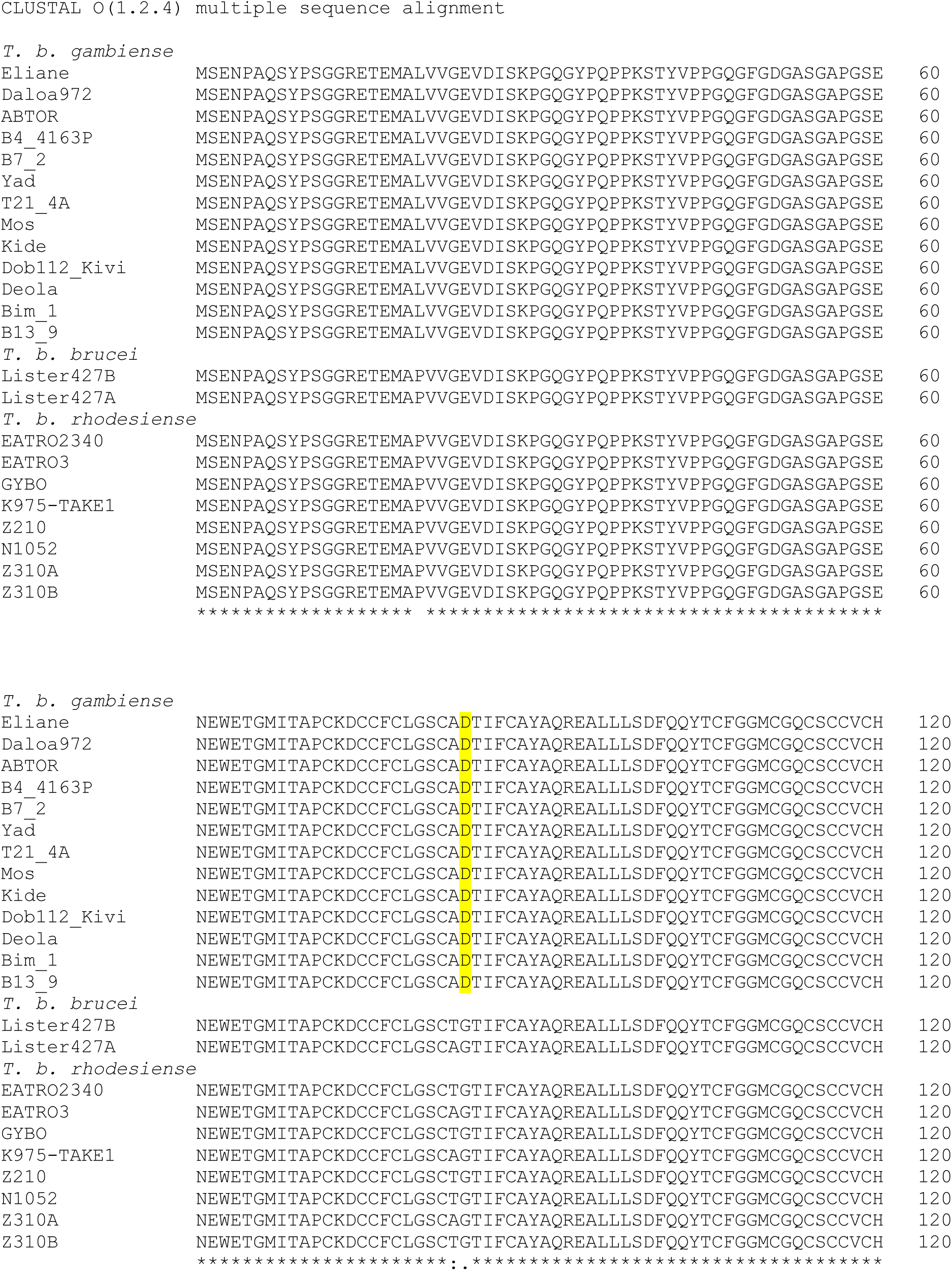
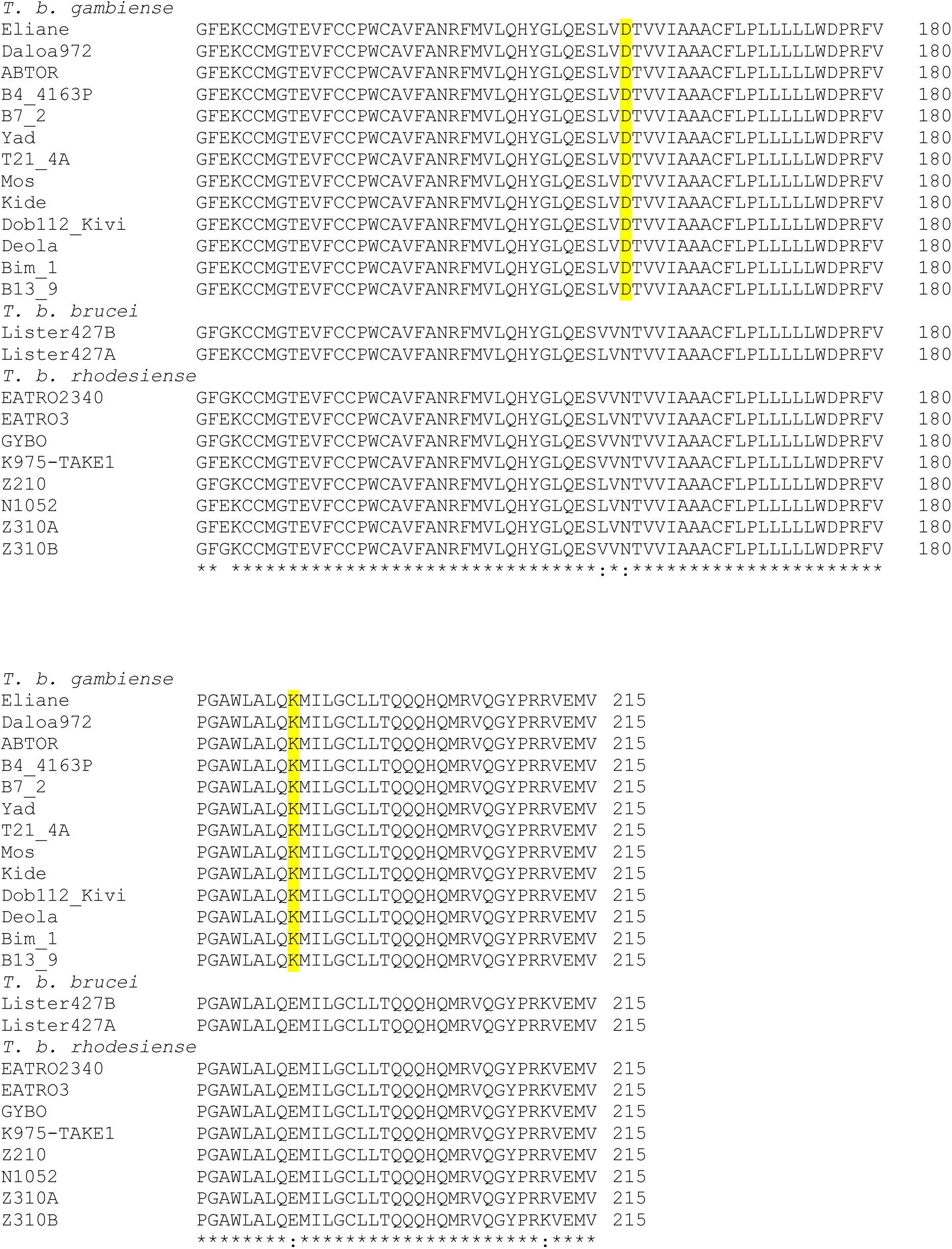

**Extended Data 1: Sequences of Tbg.972.2.1820 orthologues**

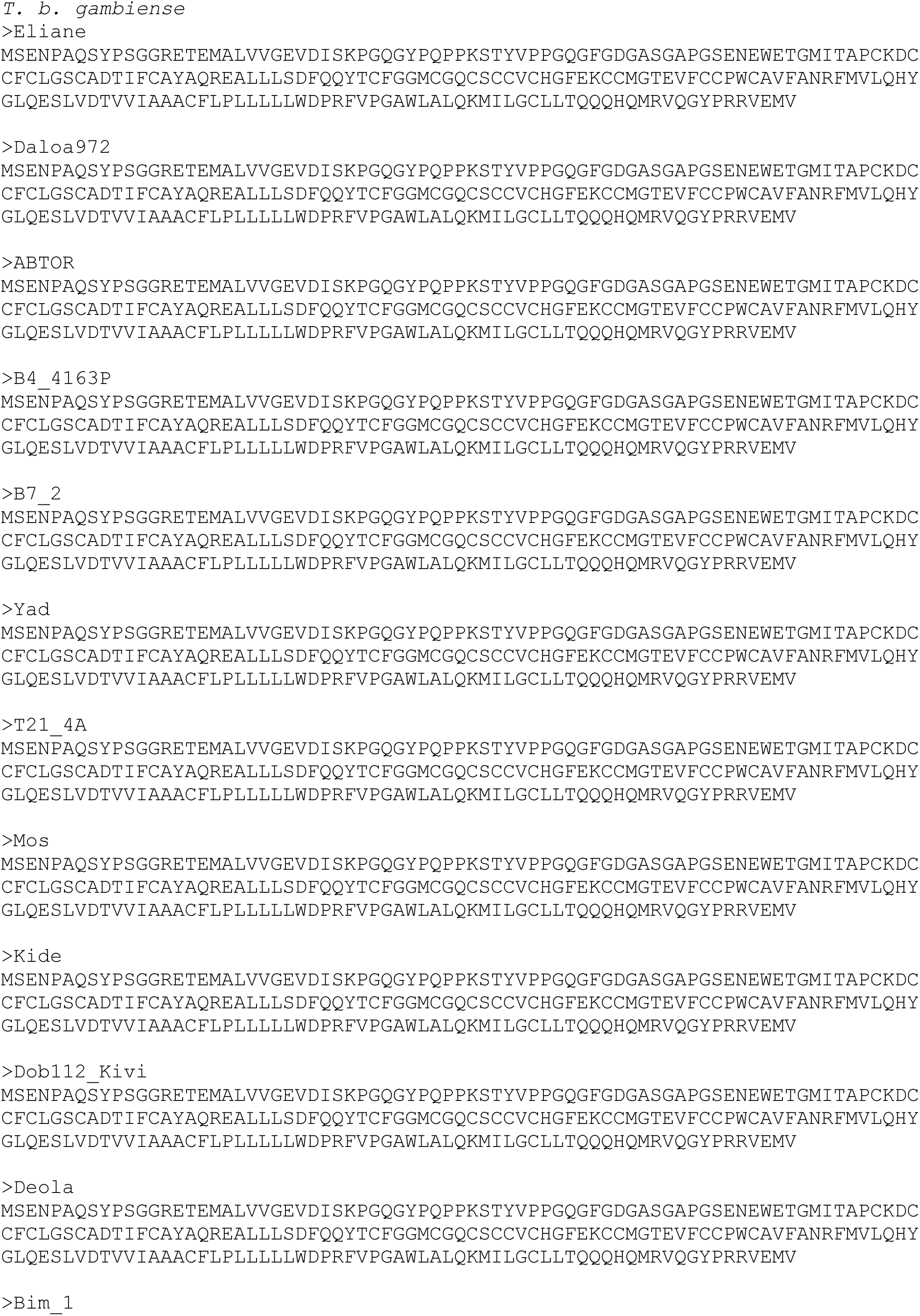
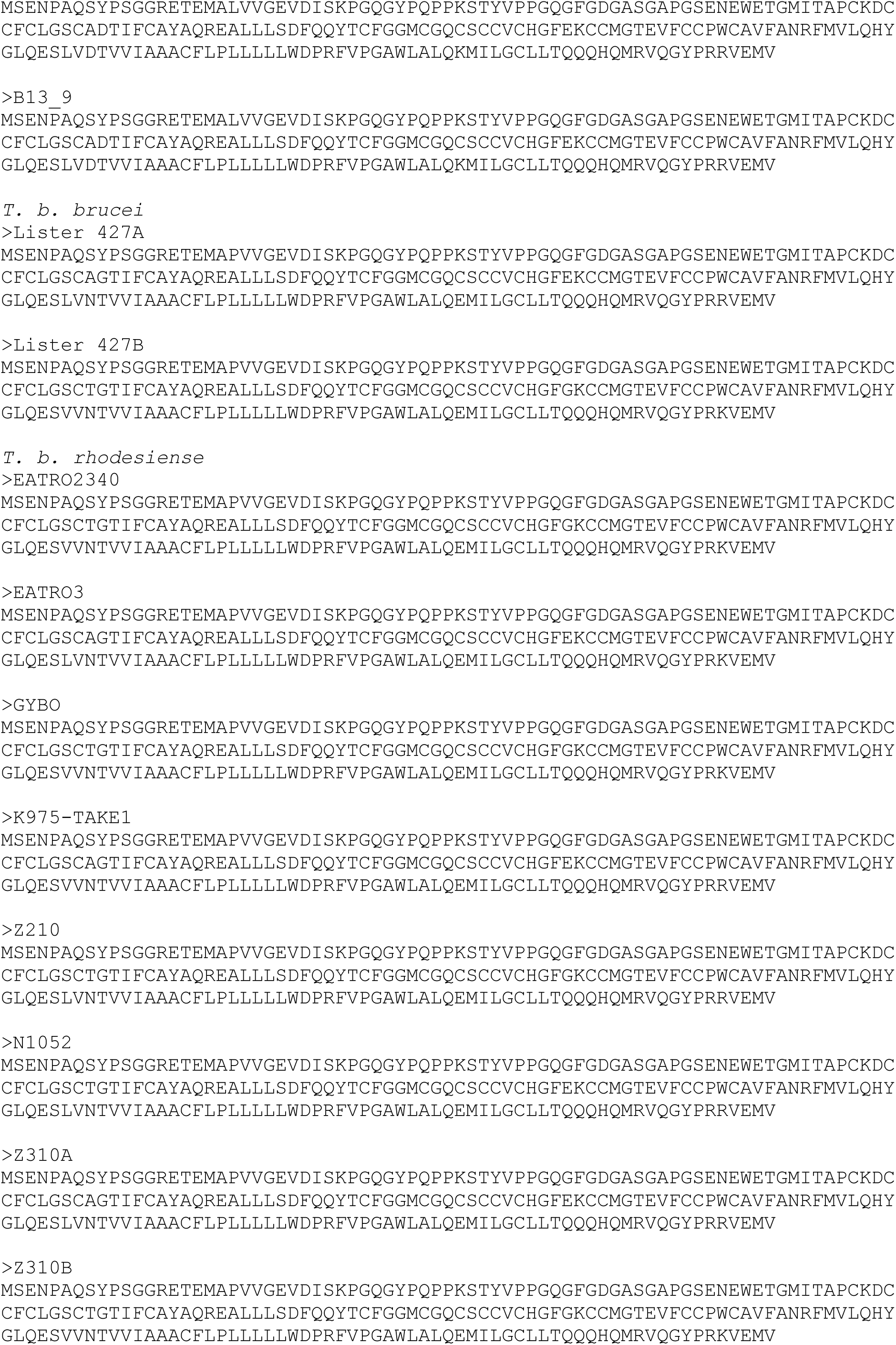

**Extended Data 2: Sequences of constructs**

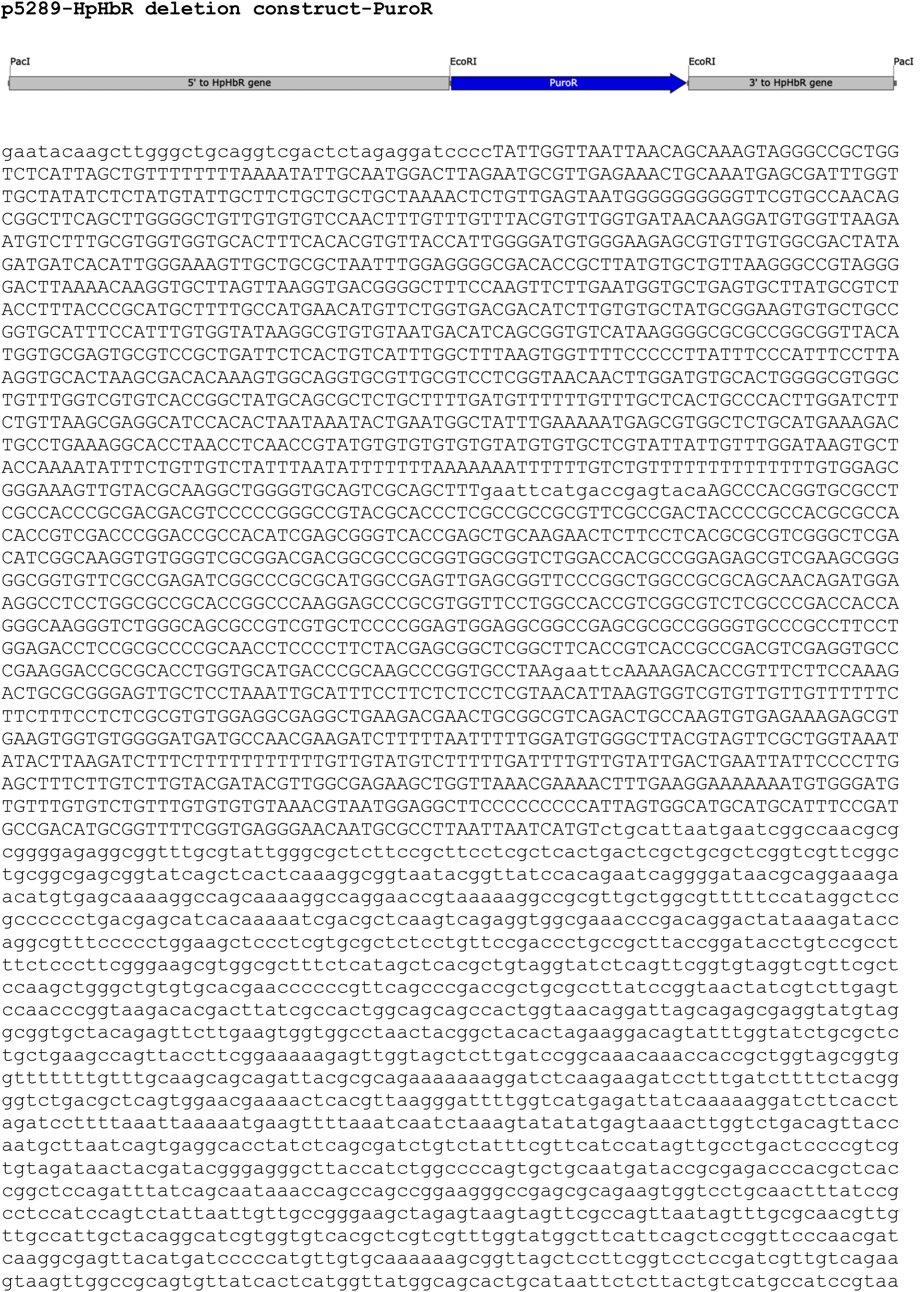
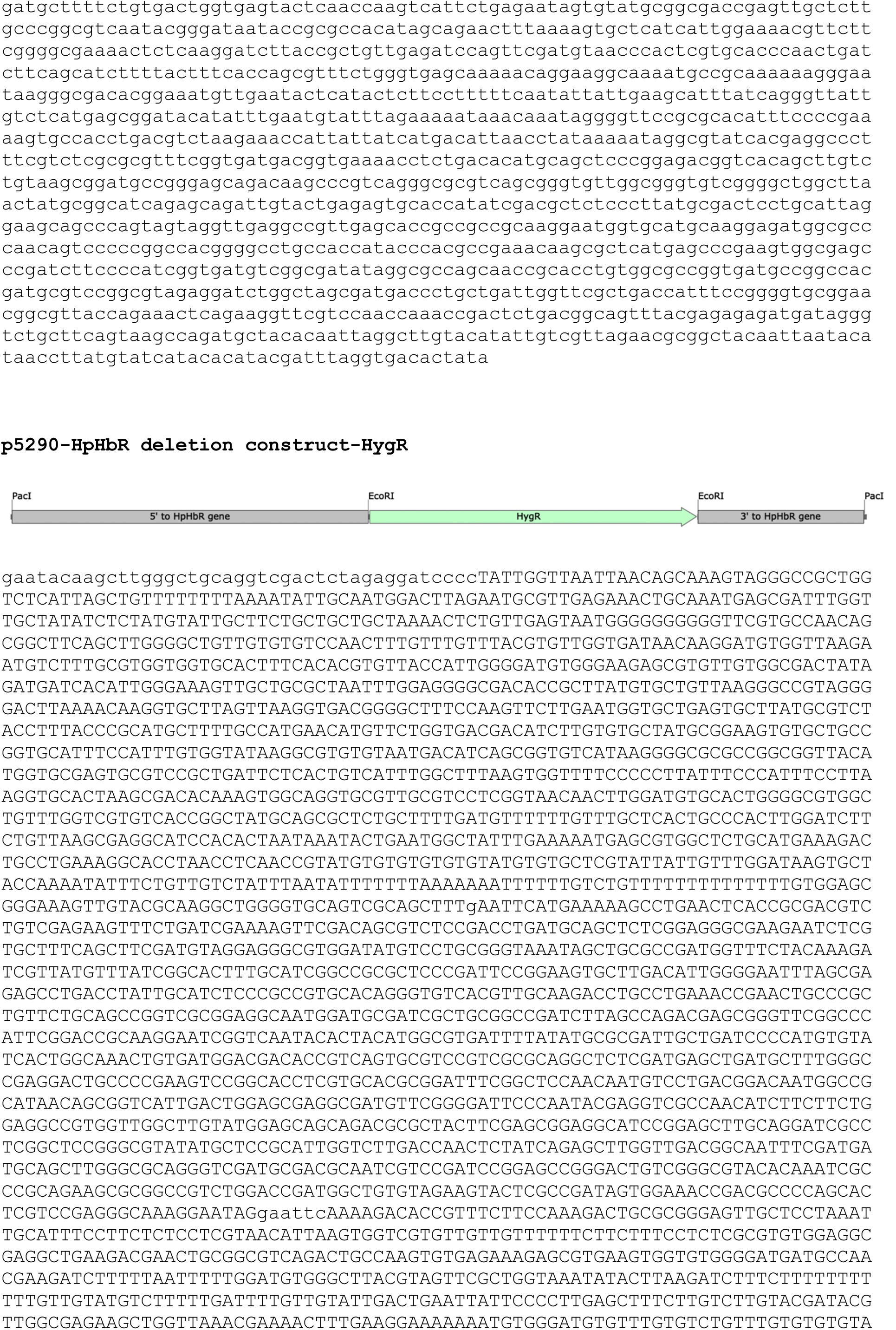
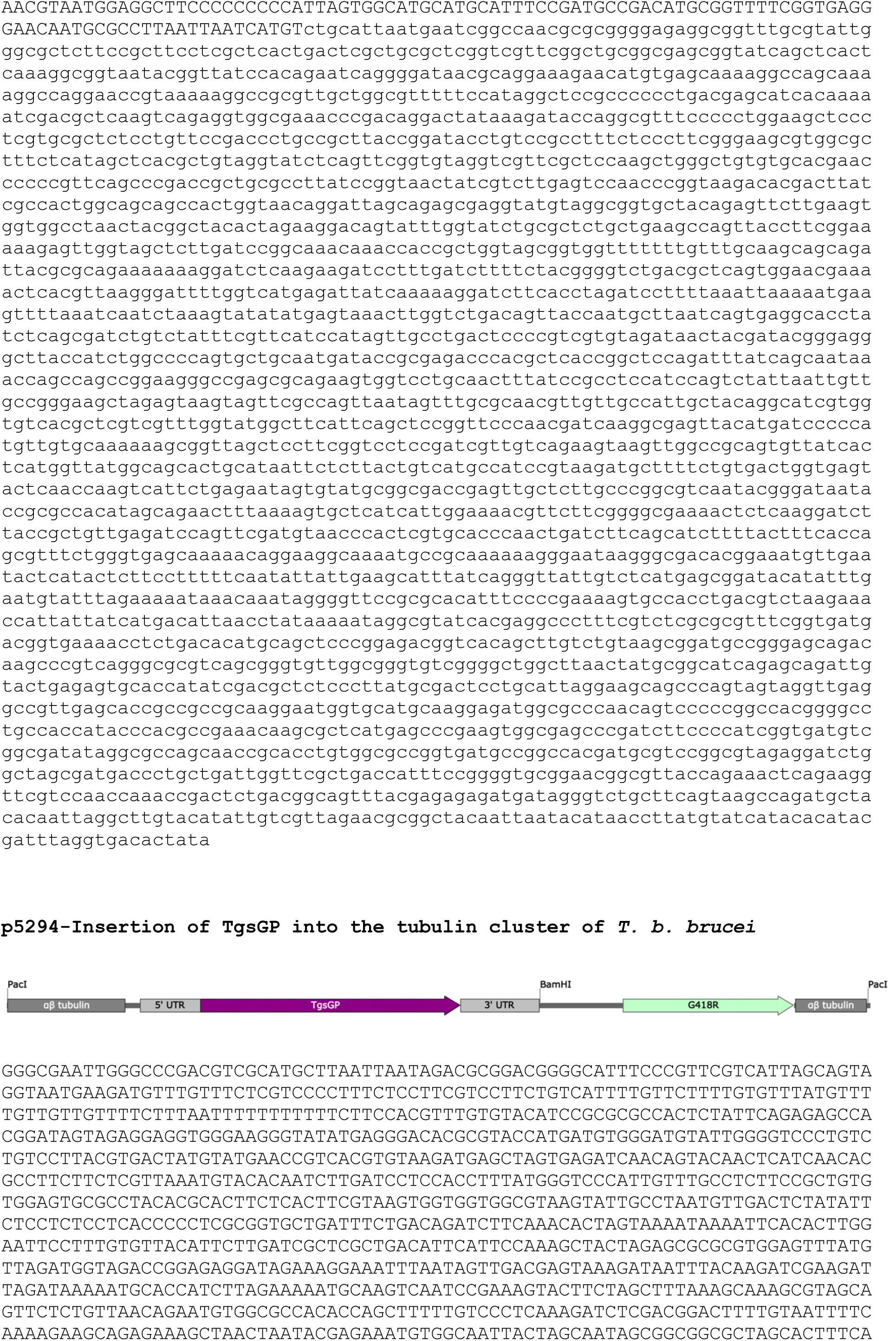
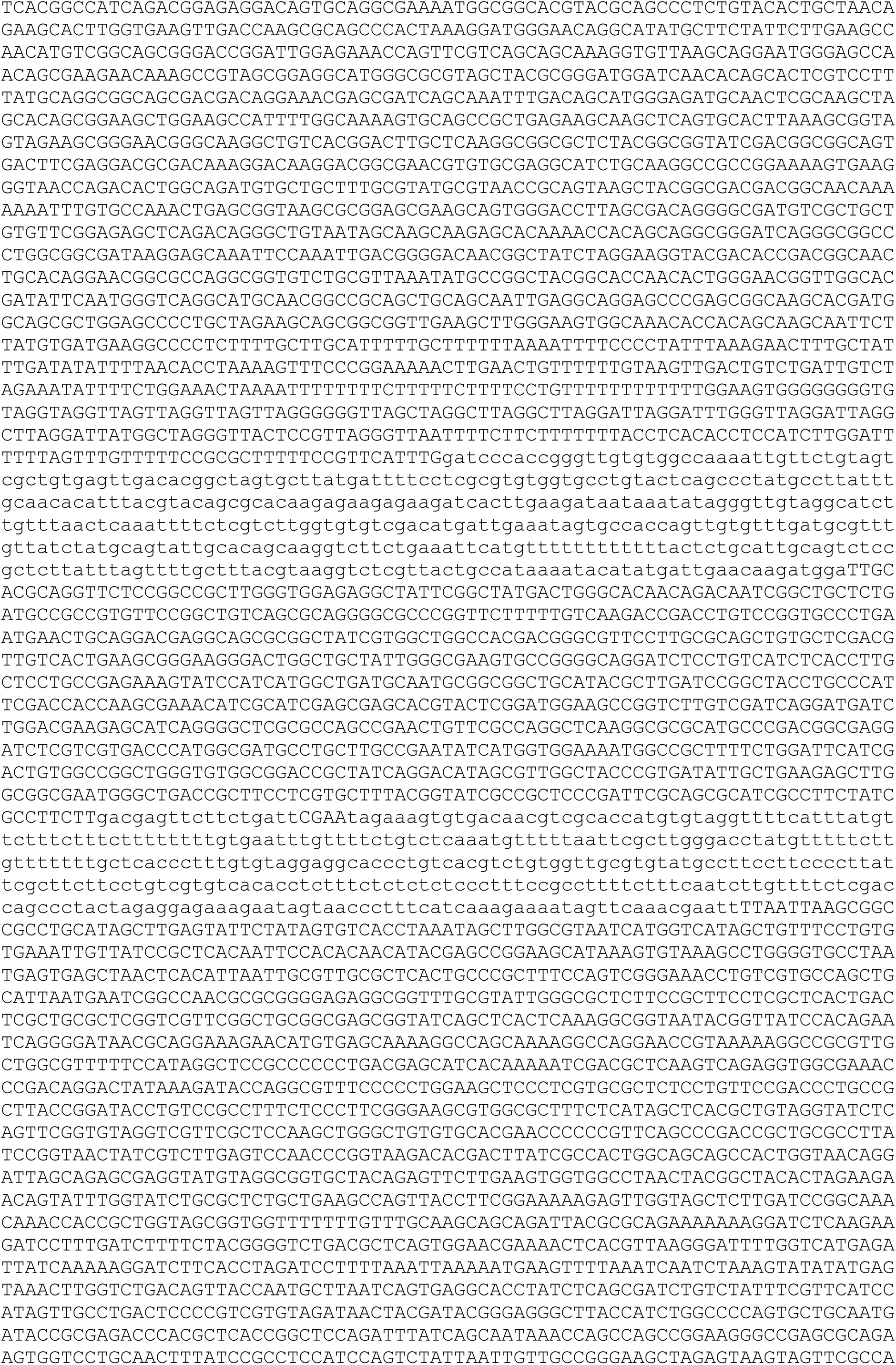
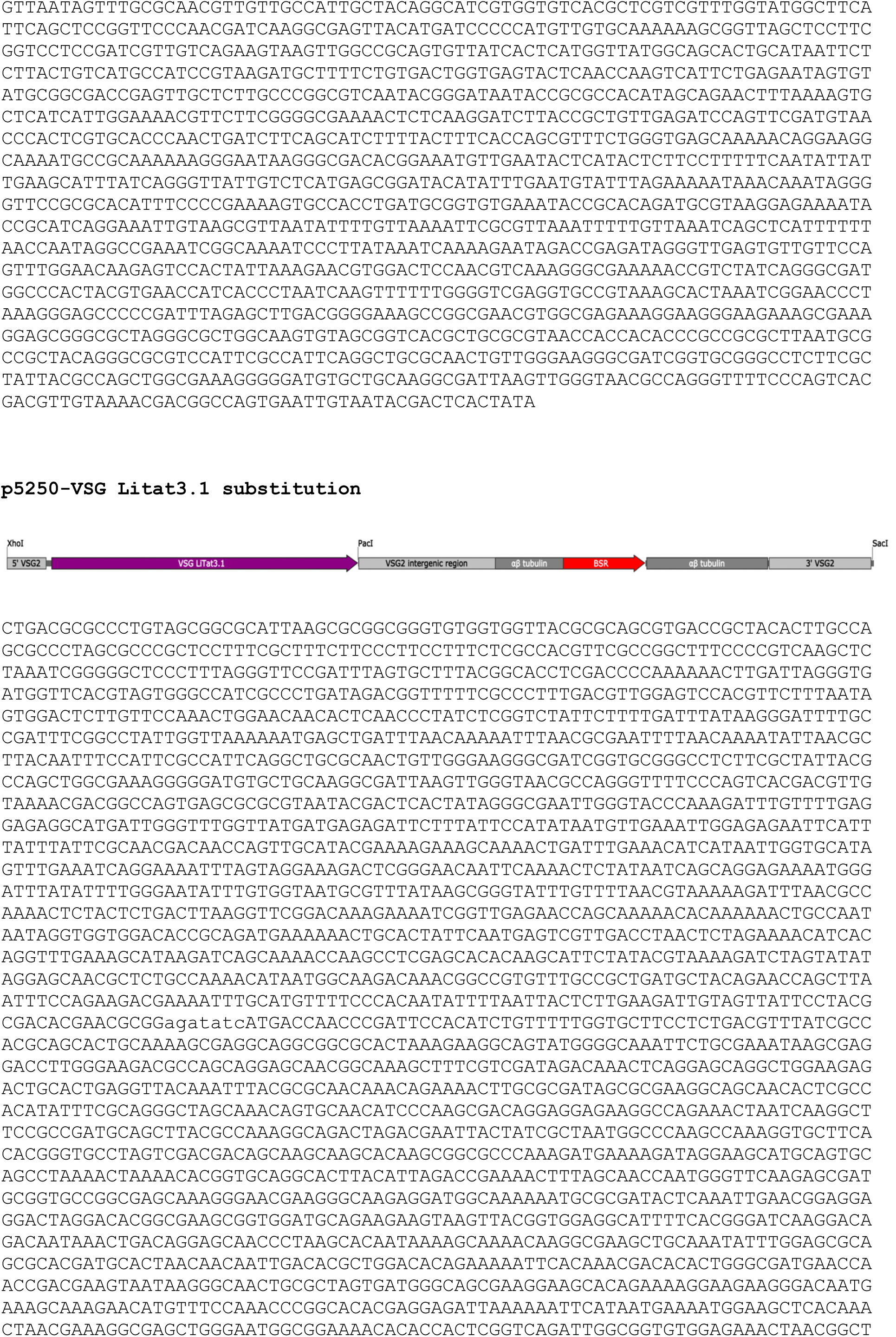
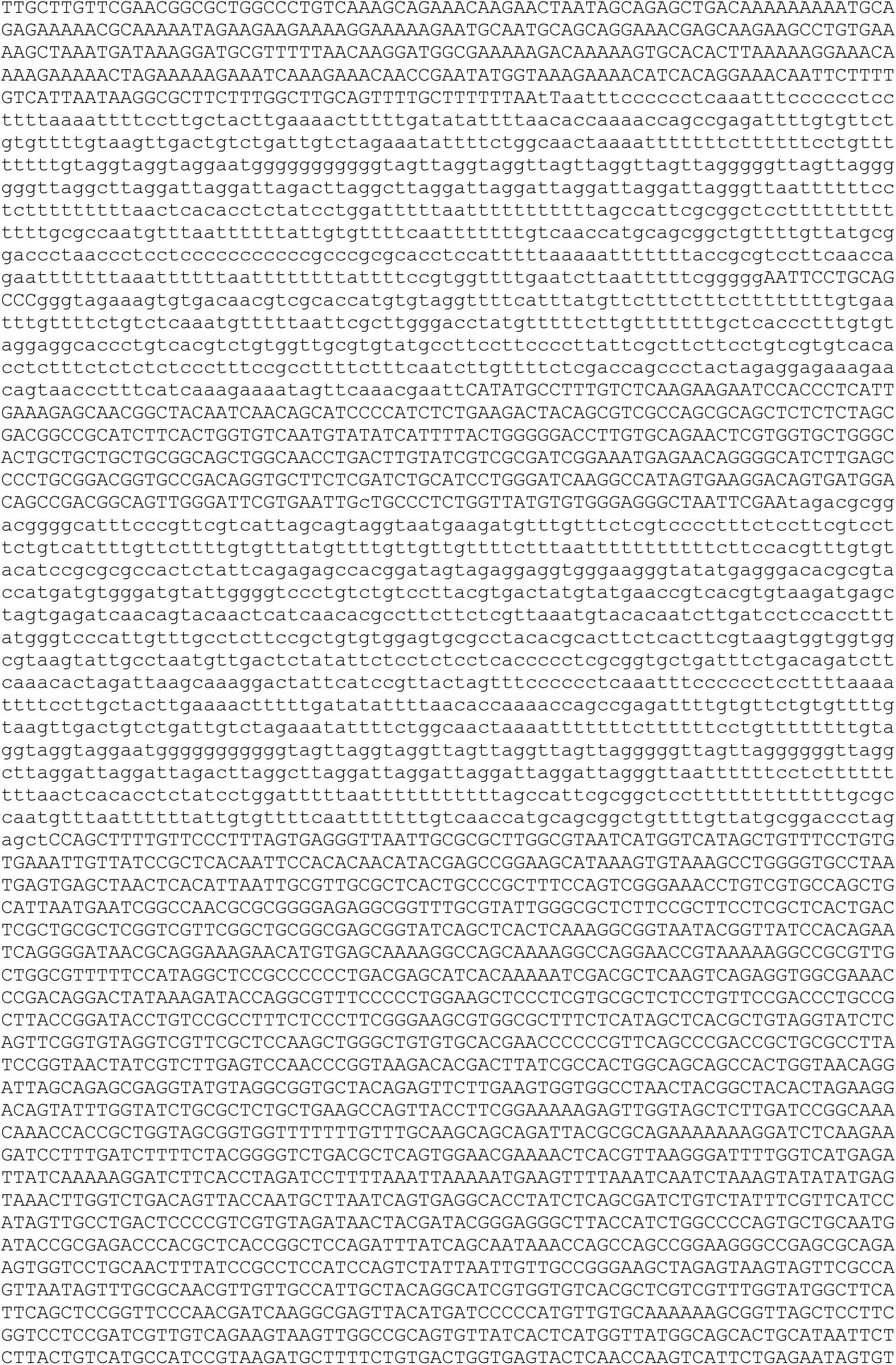
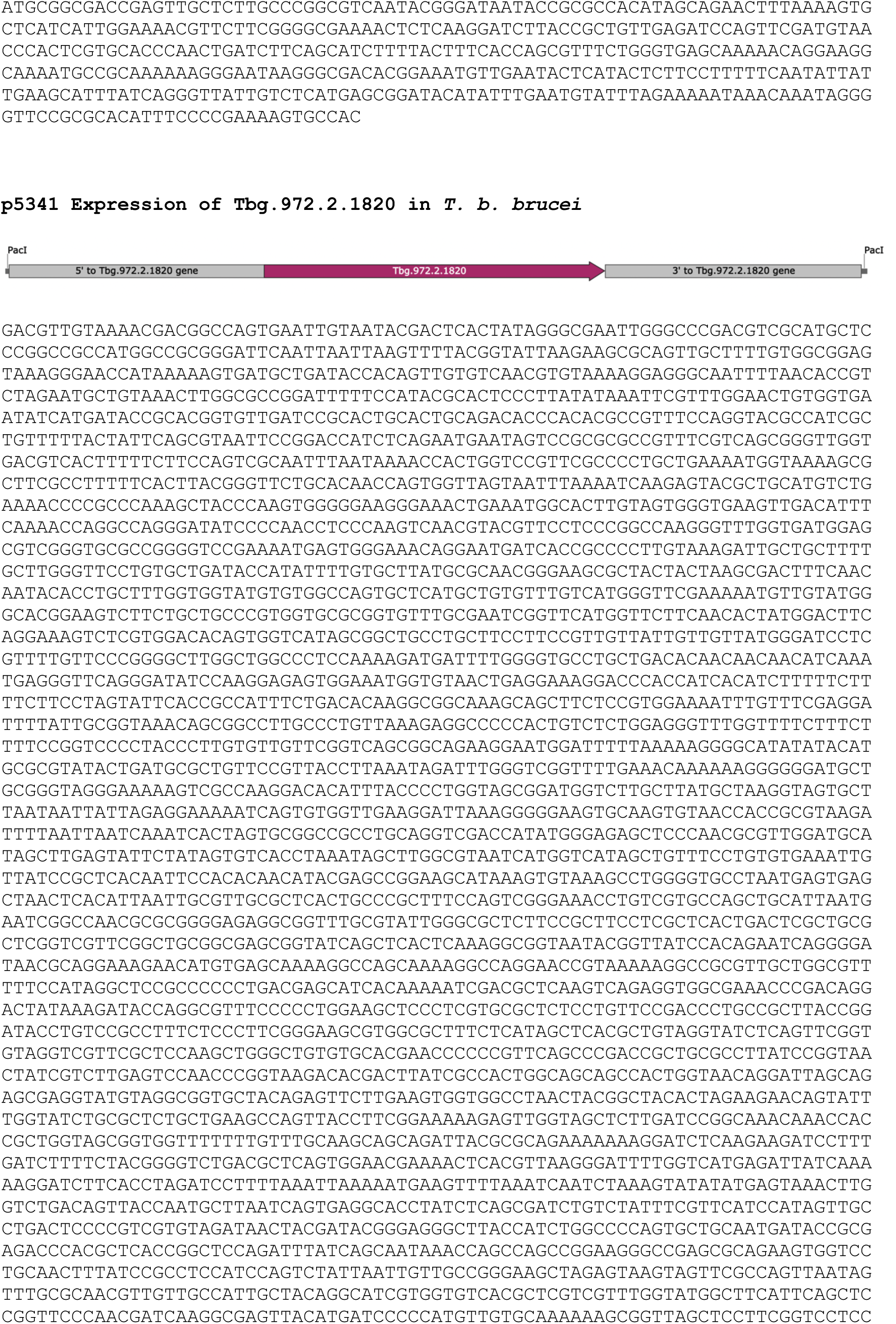
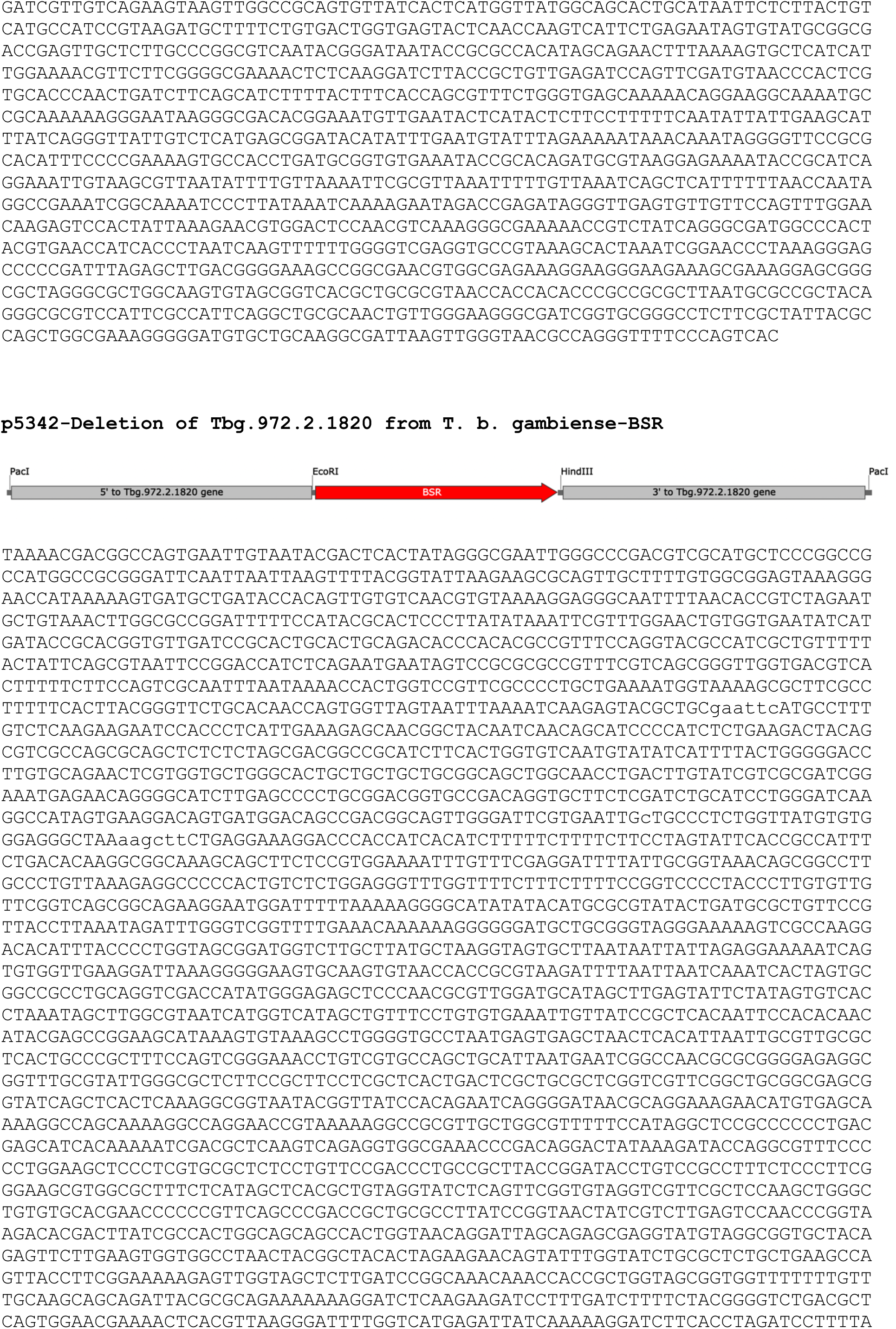
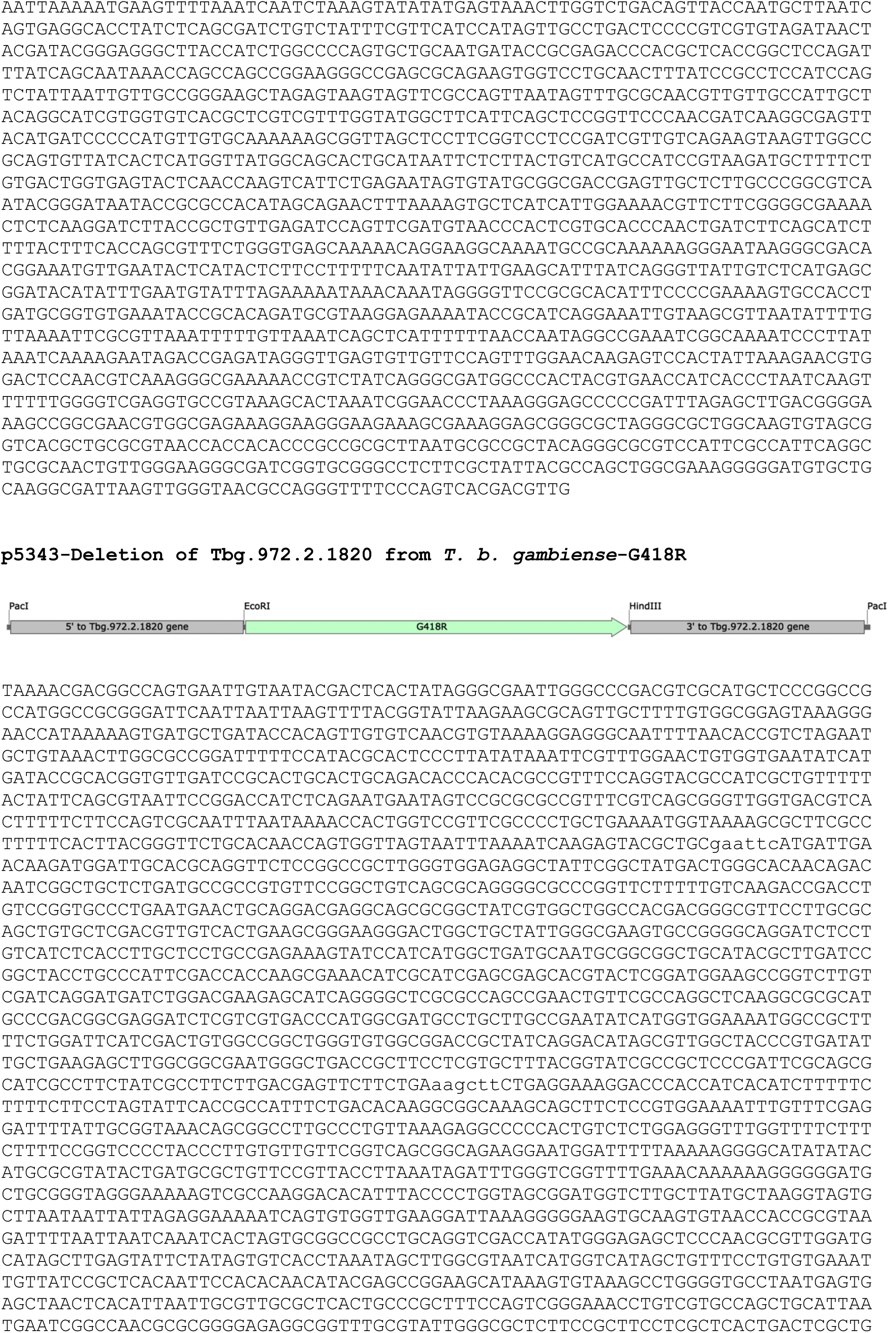
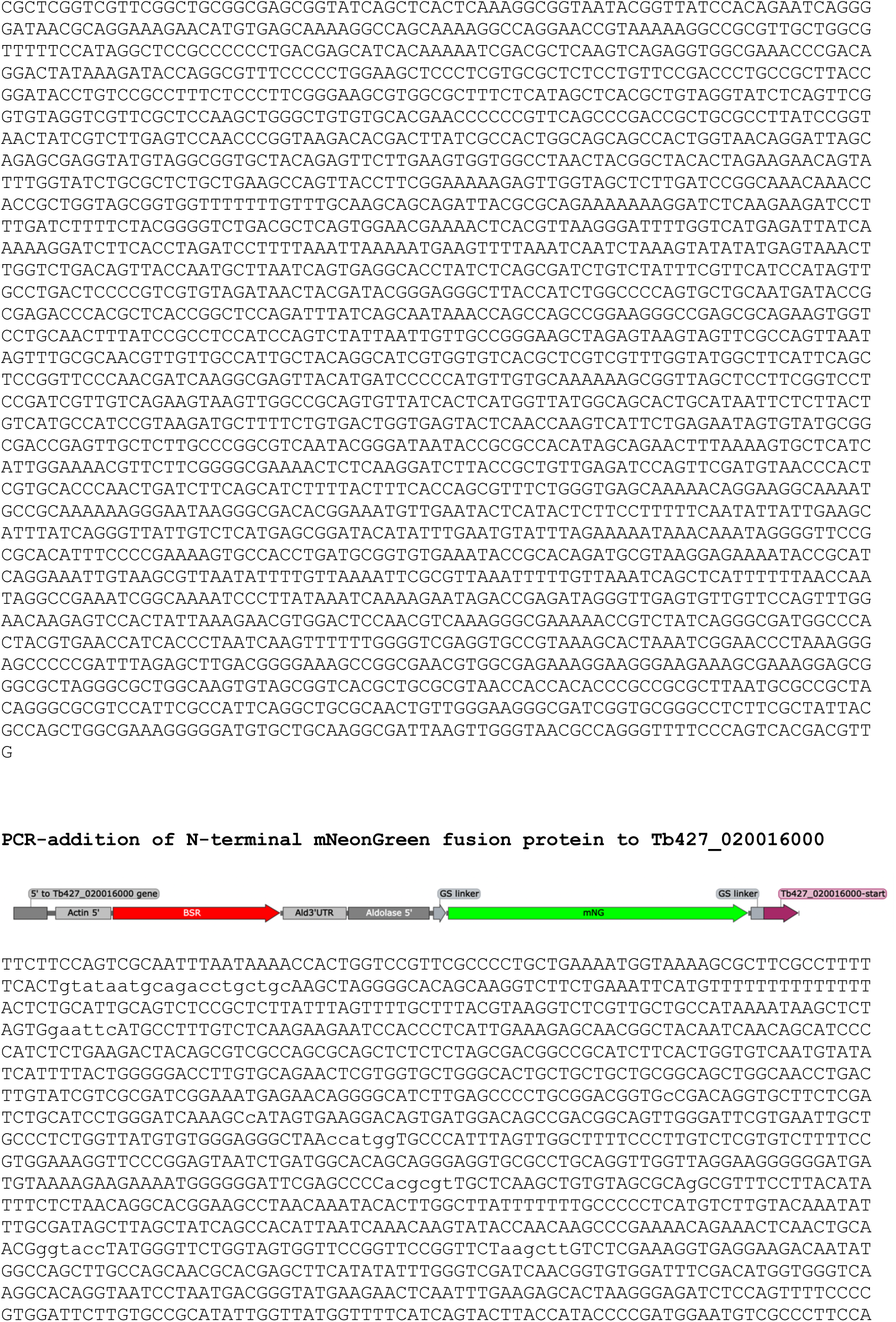
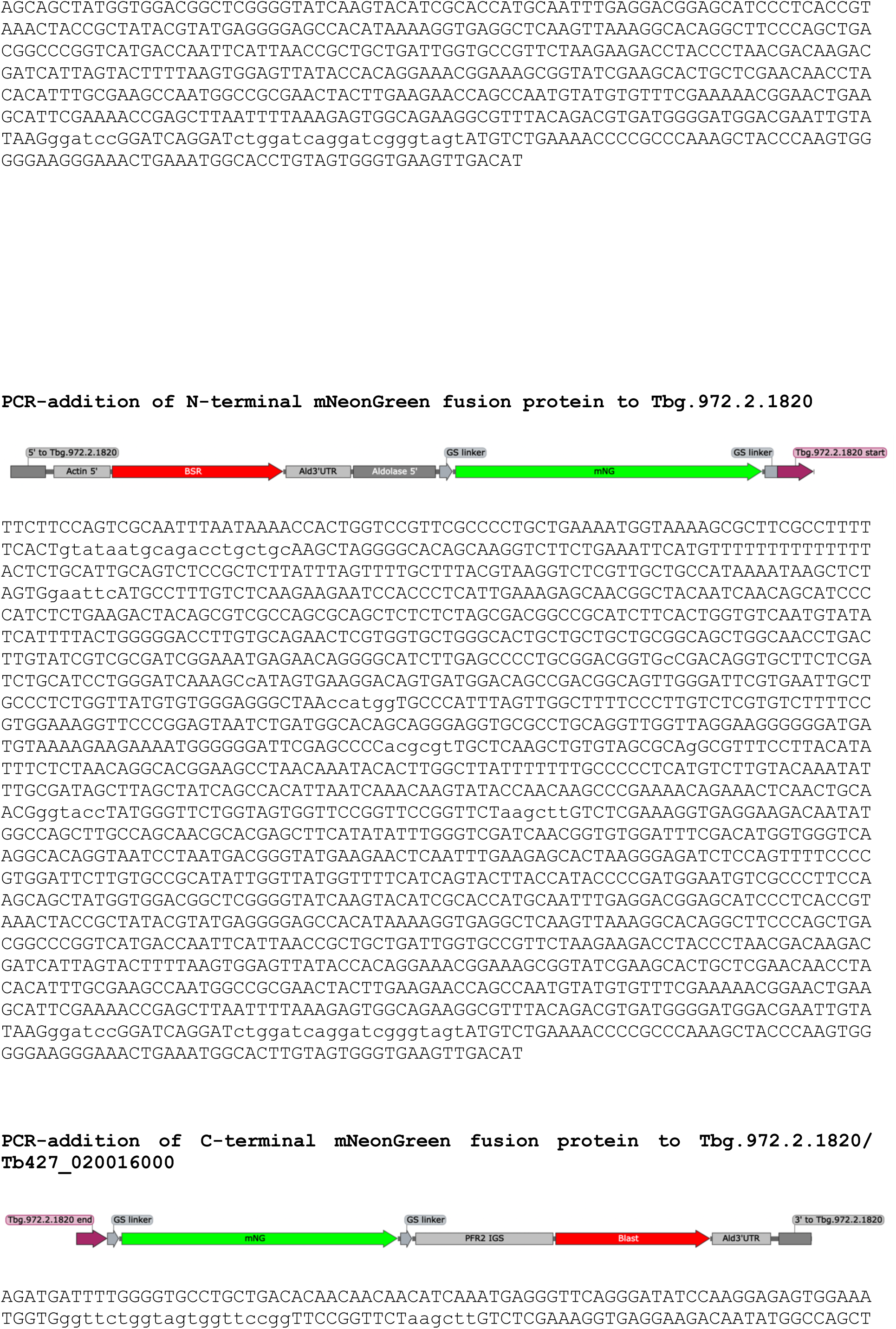
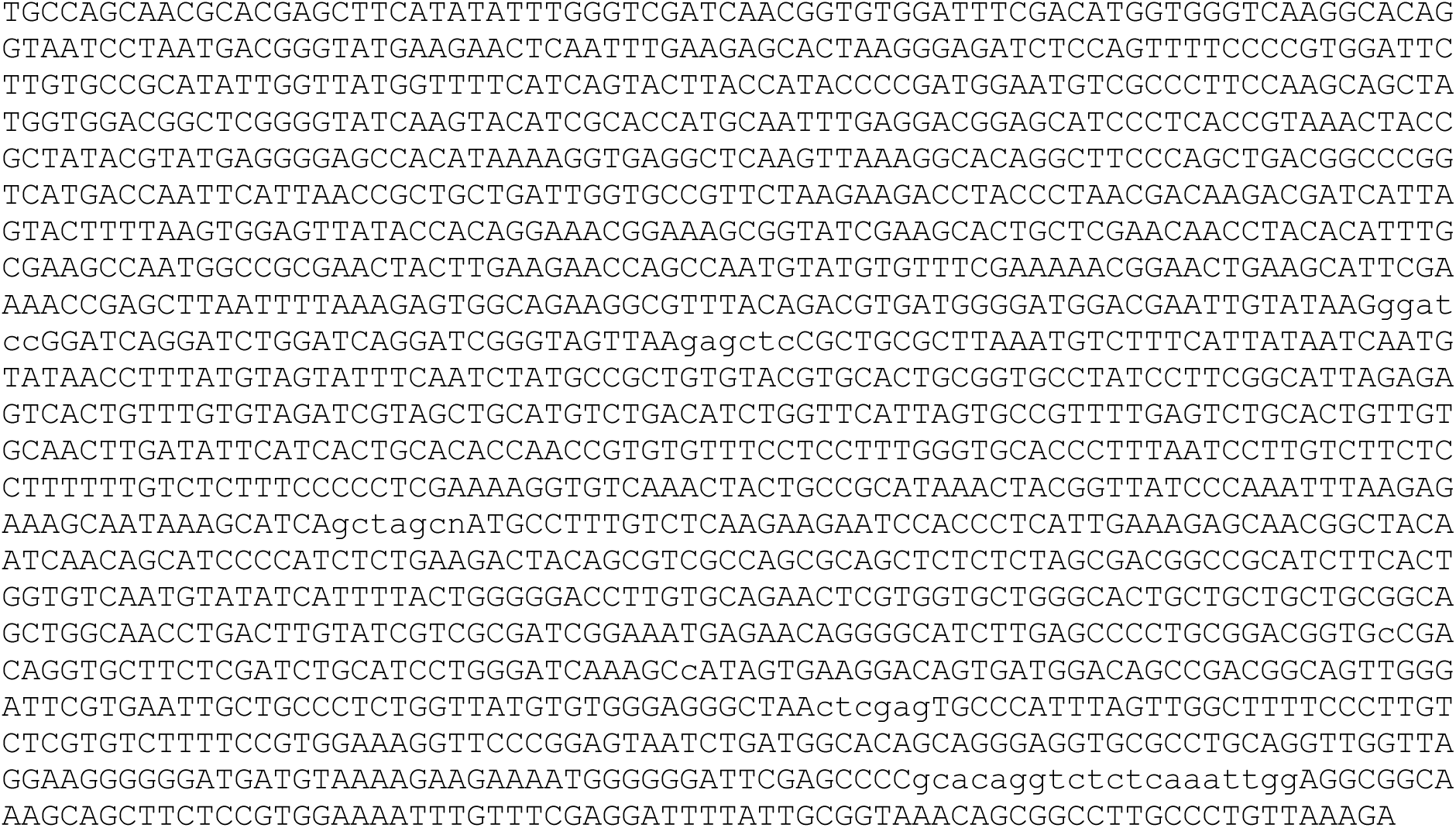

